# Combinations of low-level and high-level neural processes account for distinct patterns of context-dependent choice

**DOI:** 10.1101/508655

**Authors:** Mehran Spitmaan, Oihane Horno, Emily Chu, Alireza Soltani

## Abstract

Context effects have been explained by either low-level neural adjustments or high-level cognitive processes but not their combination. It is currently unclear how these processes interact to shape individuals’ responses to context. Here, we used a large cohort of human subjects in experiments involving choice between two or three gambles in order to study the dependence of context effects on neural adaptation and individuals’ risk attitudes. Our experiments did not provide any evidence that neural adaptation on long timescales (∼100 trials) contributes to context effects. Using *post-hoc* analyses we identified two groups of subjects with distinct patterns of responses to decoys, both of which depended on individuals’ risk aversion. Subjects in the first group exhibited strong, consistent decoy effects and became more risk averse due to decoy presentation. In contrast, subjects in the second group did not show consistent decoy effects and became more risk seeking. The degree of change in risk aversion due to decoy presentation was positively correlated with the original degrees of risk aversion. To explain these results and reveal underlying neural mechanisms, we developed new models incorporating both low- and high-level processes and used these models to fit individuals’ choice behavior. We found that observed distinct patterns of decoy effects can be explained by a combination of adjustments in neural representations and competitive weighting of reward attributes, both of which depend on risk aversion but in opposite directions. Altogether, our results demonstrate how a combination of low- and high-level processes shapes choice behavior in more naturalistic settings, modulates overall risk preference, and explains distinct behavioral phenotypes.

## Introduction

Despite the prevalent use of two-alternative choice paradigms to study value-based choice, real-life decisions often involve selecting among multiple options. The set of options, even those irrelevant to the decision maker, can strongly influence preference and alter choice processes (Tversky, 1972; Huber et al., 1982; Simonson and Tversky, 1992; Pratkanis and Farquhar, 1992; Bettman et al., 1998; Dhar and Simonson, 2003). For example, introducing a new option that is worse than one of the two existing options in all its attributes but better than the alternative option in some attributes (asymmetrically dominated option) can increase the preference for the former option (Huber et al., 1982; Dhar and Simonson, 2003; Huber and Puto, 1983; Simonson 1989). The dependence of preference on the choice set, often referred to as context or decoy effects, has been extensively studied in value-based choice and has revealed important aspects of valuation and decision processes (Bettman et al., 1998; Kahneman and Tversky, 1984; Ratneshwar et al., 1987; Payne et al., 1988; Dhar and Glazer, 1996; Wedell and Pettibone, 1996; Tsetsos et al., 2010; Trueblood, 2012).

Because of their complexity, decoy effects often have been explained by changes in subjective value due to various high-level cognitive processes, such as attentional switching to different choice attributes, menu-dependent evaluation of choice attributes, and competition between attribute processing to enhance contrast between certain attributes (Tversky, 1972; Tsetsos et al., 2010; Tversky and Simonson, 1993; Roe et al., 2001; Usher and McClelland, 2004; Johnson and Busemeyer, 2005; Usher et al., 2008; Trueblood et al., 2014). In contrast, aiming to address underlying neural mechanisms, others have attributed decoy effects to low-level adjustments of neural representations to the set of options presented on each trial (Soltani et al., 2012; Louie et al., 2013). However, none of the current models of context-dependent choice includes both low-level and high-level processes. Interestingly, long-term adaptation of neural response to the set of presented options could diminish decoy effects, and there is strong evidence for such adaptation over a block of trials or an experimental session (Louie et al., 2011; Padoa-Schioppa and Assad, 2008; Padoa-Schioppa, 2009). It is currently unknown whether such adaptation contributes to context effects or not.

In addition, although context effects mainly have been measured in a between-subjects design (Heath and Chatterjee, 1995; Pettibone and Wedell, 2000; Dhar and Sherman, 1996), there is a large variability in how such effects influence different individuals (Huber et al., 1982; Huber and Puto, 1983; Soltani et al., 2012; Pan and Lehmann, 1993; Pettibone and Wedell, 2007; Ha et al., 2009) that could reflect additional mechanisms involved in the valuation of multi-attribute options. For example, during choice between multiple gambles, the presence of a few gambles with large reward magnitudes could bias the decision maker to more strongly process reward magnitude at the expense of reward probability, ultimately resulting in more risk-seeking behavior. The influence of choice set on this selective processing of information, however, could depend on each individual’s degree of risk aversion. Currently, little is known about the dependence of context effects on risk attitudes when choosing between risky options.

Here, we measured risk preference and decoy effects in a large cohort of subjects to study whether and how context effects depend on long-term adaptation to the range of reward probabilities and magnitudes as well as on an individual’s degree of risk aversion. Risk aversion was measured using binary choice between pairs of monetary gambles (Holt and Laury, 2002). Decoy effects were measured using a phantom-decoy design; subjects were initially presented with three monetary gambles to evaluate, one of which was subsequently removed at the time of choice. To examine the contribution of neural adaptation, we introduced a new set of gambles in 20% of trials to interrupt possible ongoing long-term adaptation to the range of reward probabilities and magnitudes in the main set of gambles. Finally, we developed a new model of context-dependent choice that incorporates both low- and high-level processes in order to explain our results and reveal underlying neural mechanisms.

## Results

### Overall decoy effects

Each subject performed two experiments in which they evaluated and selected between two or three monetary gambles. In the first experiment, subjects selected between two gambles (estimation task; see Materials and Methods for more details; **Figure 1A**). Choice behavior in this experiment was used to estimate individuals’ degree of risk aversion and to tailor gambles for each subject in the second experiment. In the second experiment, subjects selected between three gambles under two different conditions (decoy task; **Figure 1B–C**). Choice behavior in this experiment was used to study context effects.

**Figure 1.**
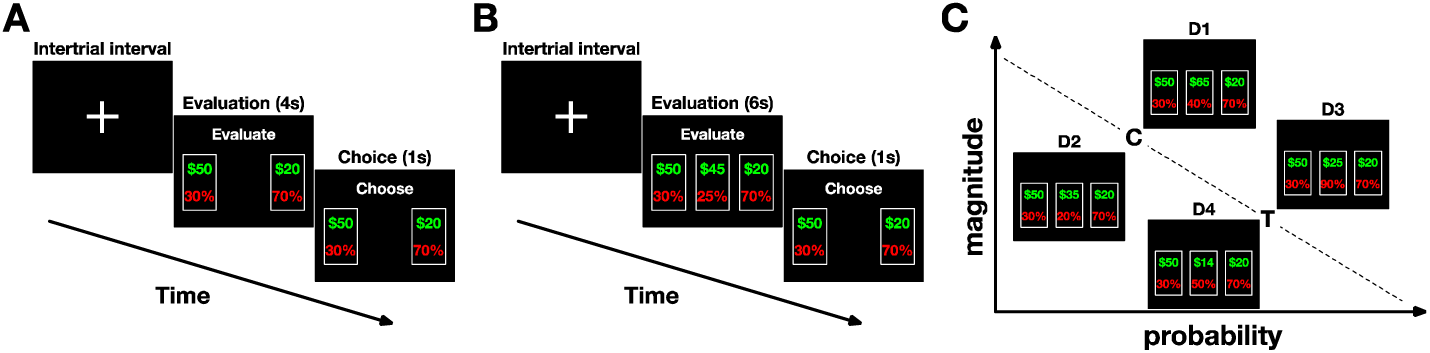
Experimental design. (**A**) Timeline of the estimation task used to measure individuals’ degree of risk aversion. After the inter-trial interval and fixation cross, two monetary gambles were presented on the screen. The subject had 4 seconds to evaluate the gambles. After “Evaluate” banner was replaced with “Choose,” the subject had 1 second to choose between the two gambles. The subjects made a choice by pressing the left or right arrow key. Selected option remained highlighted on the screen for 1 sec. (**B**) Timeline of the decoy task used to study context effects. After the intertrial interval and fixation cross, three monetary gambles were presented on the screen. The subjects had 6 seconds to evaluate the gambles. Subsequently, one of the three gambles was randomly removed, and the subjects had 1 second to choose between the remaining two gambles. The subjects made a choice by pressing the left or right arrow key; no feedback was provided following choice. (**C**) Four possible decoy positions relative to the target (T) and competitor (C) gambles are shown. For illustration purposes, decoy gambles are shown in the middle of the three gambles, but their arrangement was randomly determined on each trial of the actual experiments. D1 (D3) decoys, referred to as asymmetrically dominant decoys, were greater in both attributes than the competitor (target), whereas D2 (D4) decoys (asymmetrically dominated decoys) were worse than the competitor (target).

During the estimation task, subjects selected between two monetary gambles in each trial. One of the two gambles was always a fixed low-risk gamble: a gamble with a reward probability (*p*_*R*_) equal to 0.7 and a reward magnitude of $20. The other gamble was selected from a set of high-risk gambles with *p*_*R*_ = 0.3 and different reward magnitudes, *M*. As expected, subjects chose the high-risk gambles over the low-risk gamble more often as *M* increased (**Supplementary Figure 1A**). By fitting choice behavior with a logistic function of the difference between the two gambles’ magnitudes in a given trial, we estimated the indifference point for each subject. The indifference point was defined as the magnitude of a high-risk gamble that was as equally preferable as the low-risk gamble (see Materials and Methods). Thus, it reflects the degree of risk aversion for a given subject; a larger indifference point corresponds to increased risk aversion. Overall, we observed large variability for the indifference points across subjects (**Supplementary Figure 1B**).

During the decoy task, subjects were initially presented with three monetary gambles to evaluate, one of which was subsequently removed at the time of choice (**Figure 1B**; see Materials and Methods for more details). This phantom-decoy design allowed us to measure the influence of the removed gamble (the decoy) on the preference between the remaining gambles. Two out of the three gambles, which we refer to as the target (T) and competitor (C), were tailored for each subject using his/her indifference point estimated from the estimation task. The decoys were positioned in the attribute space relative to the target and competitor gambles in one of the four possible positions with some jitters (**Figure 1C**). In the control condition of the decoy task, jittered versions of the target, competitor, and decoy gambles were presented throughout the experiment. In the range-manipulation condition, however, we introduced a new set of gambles with large probabilities and magnitudes in 20% of trials in order to interrupt possible long-term adaptation to the range of reward probabilities and magnitudes in the main set of gambles. In total, 108 subjects participated in the study: 38 exclusively in the control condition (control only cohort), 48 exclusively in the range-manipulation condition (range-manipulation cohort), and 22 in both conditions (mixed cohort), resulting in 130 sets of data. Therefore, 60 and 70 datasets correspond to the control and range-manipulation conditions, respectively.

Examining the average probability of target selection for different decoy locations, we found a significant effect of the decoy location in the control condition (one-way ANOVA; *F*(3,236) = 6.04, *p* = 5.6 ×10^−4^; **Figure 2A**). Tukey’s *post-hoc* test revealed significant differences between the majority of decoy location pairs in terms of the average target probabilities (**Supplementary Table 1**). We also measured the effect of decoys on preference using decoy efficacies that quantify the tendency to choose the target due to the presentation of decoys at a given location after subtracting the overall tendency to choose the target (Materials and locations in the control condition (one-way ANOVA; *F*(3,236) = 31.4, *p* = 4.1 × 10^−17^; Methods). Similarly to the probability of target selection, decoy efficacies varied with decoy **Figure 2B**), and Tukey’s *post-hoc* test revealed significant differences between all pairs of decoy efficacies (**Supplementary Table 1**).

**Figure 2.**
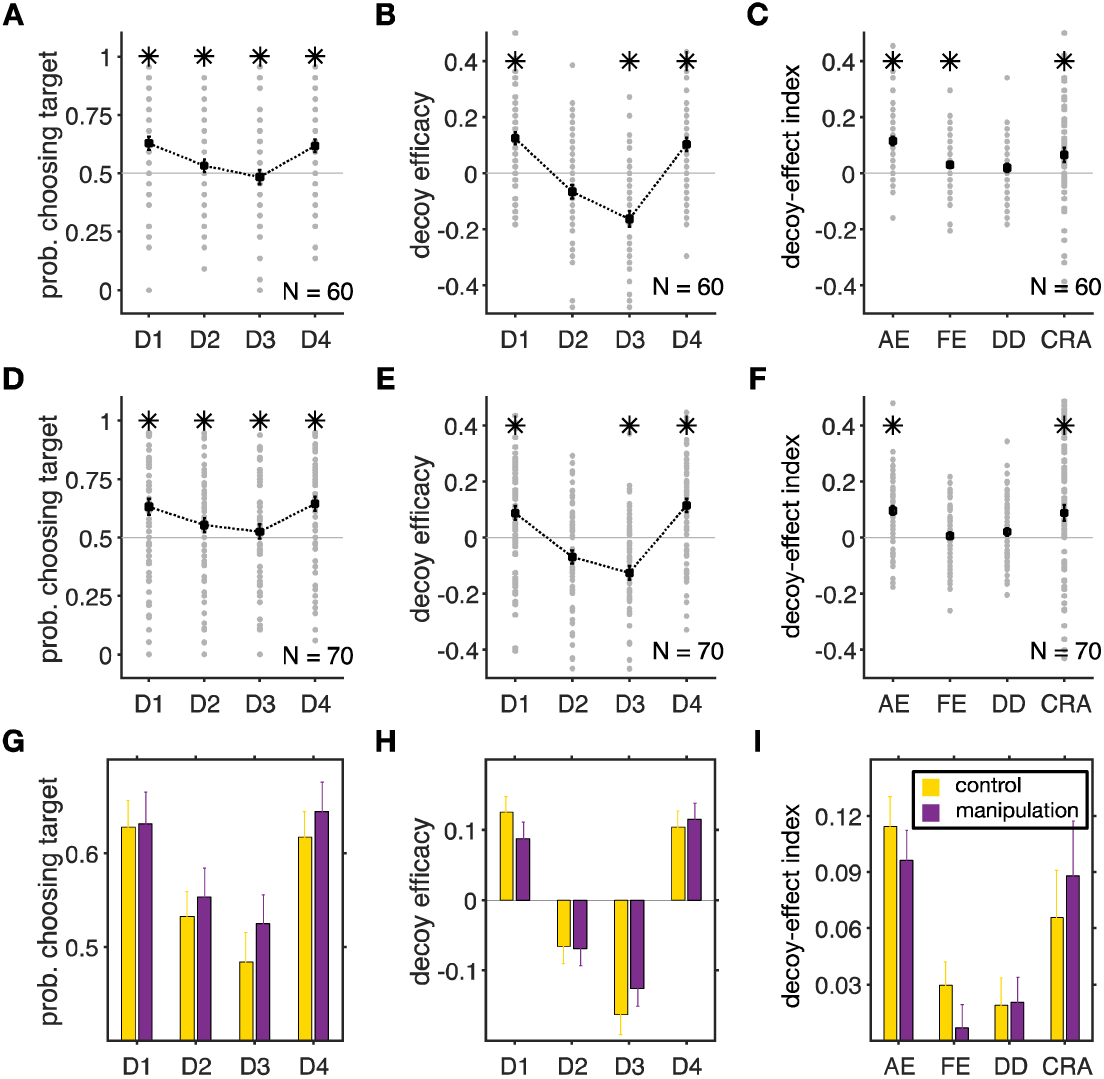
Preference was similarly influenced by decoys in the control and range-manipulation conditions. (**A**) Probability of selecting the target for different decoy locations during the control condition. Each gray circle shows the average probability that an individual subject selected the target for a given decoy location, and black squares indicate the average across all subjects. Error bars show the s.e.m., and an asterisk shows that the median of choice probability across subjects for a given decoy location is significantly different from 0.5 (two-sided Wilcoxon signed-test, *p* < 0.05). (**B**) Decoy efficacies measure the tendency to choose the target due to the presentation of decoys at a given location after subtracting the overall tendency to choose the target. An asterisk shows that the median of a given decoy efficacy across subjects is significantly different from zero (two-sided Wilcoxon signed-test, *p* < 0.05). Presentation of decoys resulted in change in preference in all locations. (**C**) Plot shows four measures for quantifying different effects of decoys on preference (AE: attraction effect; FE: frequency effect; DD: dominant vs. dominated; and CRA: change in risk aversion). Other conventions are similar to those in panel B. (**D–F**) The same as in A–C but for the range-manipulation condition. (**G–I**) Comparisons between the average target probabilities (G), decoy efficacies (H), and our four indices (I) in the two experimental conditions. Overall, there were no significant differences between the two experimental conditions based on choice probability, decoy efficacies, or any of our four measures (two-sided Wilcoxon ranksum test, *p* > 0.05).

Comparing decoy efficacies with chance level (0), we found significant effects for all decoy locations except D2. As we show later based on clustering analyses, the lack of an effect for D2 decoys could be due to a large number of certain individuals in our cohort of subjects. Asymmetrically dominant decoys decreased the preference for the gamble next to them in the attribute space (competitor for D1 and target for D3; two-sided Wilcoxon signed-test; D1: *p* = 0.023, *d* = 0.72; D3: *p* =0.042, *d* = −0.73). In contrast, asymmetrically dominated decoys increased the preference for the gamble next to them (competitor for D2 and target for D4), but this effect was only significant for D4 (two-sided Wilcoxon signed-test; D2: *p* = 0.17, *d* = −0.35; D4: *p* = 0.030, *d* = 0.59). This phenomenon is often referred to as the attraction effect (Huber et al., 1982). Considering the number of comparisons (*N* = 4) and using Bonferroni correction, most of reported decoy efficacies were not statistically significant. Finally, the decoy efficacies for the dominant and dominated decoys were anti-correlated (Pearson correlation; D1 and D3, *r* = −0.39, *p* = 0.039; D2 and D4, *r* = −0.33, *p* = 0.027), suggesting that similar mechanisms underlie their generation.

In order to fully characterize (and summarize) the changes in choice behavior due to the introduction of a decoy, we used linear transformations of decoy efficacies to define four orthogonal decoy-effect indices that measure different aspects of decoy effects in a more comprehensive way. Briefly, the overall attraction effect (AE) measures the average relative decoy effects over all locations. The overall frequency effect (FE) quantifies the overall tendency to choose the gamble next to the decoy in terms of both reward probability and magnitude (i.e., the competitor for D1 and D2 decoys, and the target for D3 and D4 decoys). The dominant vs. dominated (DD) measures the overall difference between the effects of dominant and dominated decoys. The fourth index quantifies the overall change in risk aversion (CRA) due to the decoy presentation (see Materials and Methods for more details).

Calculating these indices for each subject, we found a significant overall attraction effect (twosided Wilcoxon signed-test, *p* = 0.011, *d* = 0.94; **Figure 2C**). We also found a small but significant frequency effect (two-sided Wilcoxon signed-test, *p* = 0.046, *d* = 0.31), indicating an overall increase in the selection of the gamble next to the decoys. However, considering the number of comparisons (*N* = 4) and using Bonferroni correction, the frequency effect was not statistically significant in the control condition. There was no significant difference in the effects of dominant and dominated decoys as measured by the DD (two-sided Wilcoxon signed-test, *p* = 0.22, *d* = 0.17). Finally, because of the symmetry in decoy presentation as well as the tailoring of the target and competitor gambles for individual subjects, each subject should choose equally between the target and competitor in absence of any decoy effects. Therefore, any overall bias toward either gamble could be attributed to changes in risk preference due to the presentation of decoys, which is quantified by the CRA. Interestingly, we found a significant increase in risk aversion due to decoys across all subjects (two-sided Wilcoxon signed-test, *p* = 0.035, *d* = 0.33). Again, considering the number of comparisons (*N* = 4) and using Bonferroni correction, this effect was not statistically significant. However, as we show below this is mainly due to the presence of two distinct phenotypes of subjects that change their overall risk preference in opposite directions.

### Contribution of long-term neural adaptation to decoy effects

Previous monkey experiments from Padoa-Schioppa’s group (Padoa-Schioppa, 2009) have shown adaptations of neural response to the range of stimuli on the order of 100 trials. In order to test the contribution of such long-term adaptation to context effects, we introduced a new set of gambles with large reward probabilities and magnitudes during 20% of trials of the range-manipulation condition (see Materials and Methods). We predicted that such adaptation should decrease decoy effects. In an extreme case, full adaptation to the range of presented gambles could diminish decoy effects because the range of all stimuli could be incorporated into representations of reward attributes and thus, no range normalization would be necessary. We set the attributes of high-range gambles as large as possible (considering that we had to resolve such gambles if they were chosen) to increase the chance of detecting an effect.

Computing the average probability of target selection for different decoy locations, we found a significant effect of the decoy location in the range-manipulation condition (one-way ANOVA; *F*(3,236) = 3.45, *p* = 0.017; **Figure 2D**). Tukey’s *post-hoc* test revealed significant differences between all decoy location pairs in terms of the average target probabilities (**Supplementary Table 1**). Similarly to the probability of target selection, decoy efficacies varied with decoy locations (one-way ANOVA; *F*(3,236) = 24.3, *p* = 5.9 ×10^−14^; **Figure 2E**), and Tukey’s. *post-hoc* test revealed significant differences between all pairs of decoy efficacies (**Supplementary Table 1**).

Similar to the control condition, subjects in the range-manipulation condition exhibited significant decoy effects for D1, D3, and D4 decoys (two-sided Wilcoxon signed-test; D1: *p =* 0.039, *d* = 0.43; D2: *p =* 0.33, *d =* −0.34; D3: *p =*0.031, *d =* −0.62; D4: *p =* 0.028, *d =* 0.59; **Figure 2E**). In addition, we found a significant overall attraction effect and change in risk aversion due to decoy presentation (two-sided Wilcoxon signed-test; AE: *p* = 0.014, *d* = 0.72; FE: *p* = 0.61, *d* = 0.07; DD: *p* = 0.73, *d* = 0.18; CRA: *p* = 0.044, *d* = 0.36; **Figure 2F**). Considering the number of comparisons (*N* = 4) and using Bonferroni correction, however, most of reported decoy efficacies were not statistically significant. Finally, similar to the control condition, the decoy efficacies for the dominant and dominated decoys were anti-correlated in the range-manipulation condition (Pearson correlation; D1 and D3, *r* = −0.38, *p* = 0.002; D2 and D4, *r* = −0.37, *p* = 0.001).

We next directly compared decoy efficacies and decoy-effect indices across the control and range-manipulation conditions. Despite the large number of subjects in our experiment, however, we did not find any significant difference between subjects’ behavior in the two experimental conditions with regards to the average decoy efficacies (two-sided Wilcoxon ranksum test; D1: *p* = 0.51; D2: *p* = 0.92; D3: *p* = 0.35; D4: *p* = 0.68; **Figure 2H**) or decoy-effect indices (two-sided Wilcoxon ranksum test; AE: *p* = 0.31; FE: *p* = 0.11; DD: *p* = 0.42; CRA: *p* = 0.26; **Figure 2I**). These results provide no evidence that range manipulation, designed to interrupt long-term adaptation to the set of options, has an influence on decoy effects.

To test for interactions between experimental conditions and decoy effects, we also performed two-way ANOVA using both experimental conditions (control and range manipulation). We found that the decoy location had significant effects on average target probabilities (two-way unequal size ANOVA; *F*(3,512) = 8.80, *p* = 0) and decoy efficacies (two-way unequal size ANOVA; *F*(3,512) = 54.67, *p* = 0). However, we did not find any effect for experimental conditions or their interactions with decoy locations on average target probabilities (two-way unequal size ANOVA; experimental conditions: *F*(1,512) = 1.15, *p* = 0.28; interaction: *F*(3,512) = 0.13, *p* = 0.94) or on decoy efficacies (two-way unequal size ANOVA; experimental conditions: *F*(1,512) = 0.01, *p* = 0.93 interaction: *F*(3,512) = 0.82, *p* = 0.48 **Figure 2 G,H**).

Combining data from the control and range-manipulation conditions, we found significant decoy efficacies for all decoy locations except D2, even after considering Bonferroni correction (twosided Wilcoxon signed-test; D1: *p* = 0.005, *d* = 0.68; D2: *p* = 0.15, *d* = −0.21; D3: *p* =0.001, *d* = - 0.65; D4: *p =*0.004, *d =* 0.62). In addition, we found a significant overall attraction effect and change in risk aversion due to decoy presentation (two-sided Wilcoxon signed-test; AE: *p* = 4.8 × 10^−12^, *d* = 1.34 FE: *p* = 0.08, *d* = 0.23; DD: *p* = 0.78, *d* = 0.06; CRA: *p* = 3.86 × 10^−4^, *d* = 1.18).

We also examined the time course of decoy-effect indices in the two experimental conditions by computing the running average of AE and CRA over time, using a sliding window with a length of 40 trials and steps of 20 trials (**Figure 3**). However, we did not observe any significant changes in any of the decoy-effect indices over time (two-sided t-test; AE (control), *p* = 0.96; AE (range-manipulation), *p* = 0.75; CRA (control), *p* = 0.81; CRA (range-manipulation), *p* = 0.46). Finally, there was no significant differences in the slopes of the AE or CRA as a function of time (within a session) between the two experimental conditions (AE (control), *b* = 0.0004, 95% CI [- 0.0146, 0.0154]; AE (range-manipulation), *b* = −0.0027, 95% CI [−0.0191, 0.0137]; CRA (control), *b* = 0.0021, 95% CI [−0.0149, 0.0191]; CRA (range-manipulation), *b* = 0.0052, 95% CI [−0.0144, 0.0248]).

**Figure 3.**
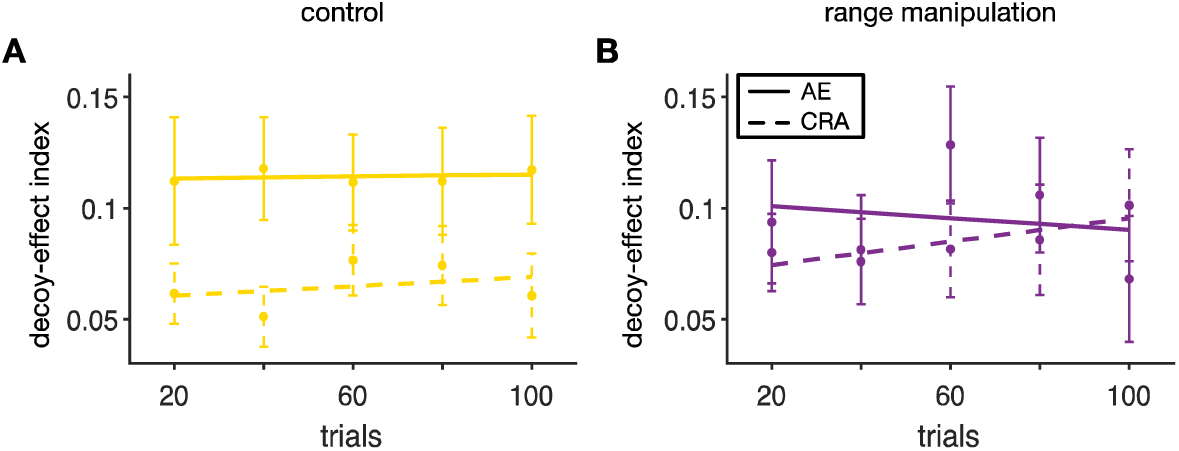
No significant difference between the time course of decoy effects in the control and range-manipulation conditions. Plotted is time course of the overall attraction effect (AE) and the overall change in risk aversion (CRA) in the control (**A**) and range-manipulation conditions (**B**). Each point shows the average value of the respective measure during 40-trial bins around that point across all subjects. The error bar shows the s.e.m. The solid and dashed lines show the regression lines.

Together, our analyses did not provide any evidence for changes in decoy effects due to range manipulation, which was intended to interrupt possible long-term adaptation to the range of reward probabilities and magnitudes. Therefore, we did not find any evidence that long-term neural adaptation on the order of ∼100 trials––present in the control but not in the rangemanipulation condition––contributes to context effects.

### Distinct behavioral phenotypes for susceptibility to decoy

Our data analyses above did not reveal any evidence for the effect of range manipulation on context effects, which could be due to many reasons including ineffectiveness of our manipulation, individual variability, etc. Nonetheless, the control and range-manipulation conditions provided us with a large dataset to investigate individual variability and behavioral phenotype with respect to decoy effects. To that end, we used a strict clustering method to isolate distinct patterns of decoy effects in our large cohort of subjects. For this whole-data clustering analysis, we used all 130 datasets. Nevertheless, we performed additional analyses to ensure that there was no difference between the three cohorts of subjects in terms of behavioral phenotypes identified via clustering (see below). We adopted the k-means clustering method for different sets of features and numbers of clusters. The sets of features included all the possible exclusive combinations of four decoy-effect indices and the cluster sizes ranged from 2 to 5. We found the best clustering (based on silhouette values) is achieved using all decoy-effect indices and two clusters (**Supplementary Figure 2**). We used the clusters with the best separation to divide the subjects into two groups (Group 1: N = 72, and Group 2: N = 58).

The subjects in the two groups exhibited very distinct patterns of responses to the presentation of decoys (**Figure 4**). More specifically, subjects in Group 1 showed an overall increase in risk aversion during the decoy task by selecting the target with a probability larger than 0.5 (over all decoy locations), whereas subjects in the Group 2 showed a decrease in risk aversion by choosing the target with a probability smaller than 0.5 (**Figure 4A–B**). In addition, subjects in the first group showed strong and consistent (i.e. across all decoy types) decoy effects (two-sided Wilcoxon ranksum test; D1: *p* = 0.001, *d* = 0.99; D2: *p* = 0.008, *d* = −0.62; D3: *p* = 0.0001, *d* = - 0.67; D4: *p* = 0.0003, *d* = 0.60; **Figure 4C**), whereas decoy effects in the second group were inconsistent and limited to decoys next to the less risky gamble (two-sided Wilcoxon ranksum test; D1: *p* = 0.42, *d* = 0.18; D2: *p* = 0.79, *d* = −0.03; D3: *p* = 0.0001, *d* = −0.67; D4: *p* = 0.0022, *d* = 0.58; **Figure 4D**). Specifically, the lack of an effect for D2 decoys in Group 2 could explain the absence of this effect in overall data. That is, the absence of significant decoy effect for decoys in D2 location compared to a previous study by Soltani et al. (2012) could be the result of having more subjects with inconsistent decoy effects (Group 2 subjects) in the current study.

**Figure 4.**
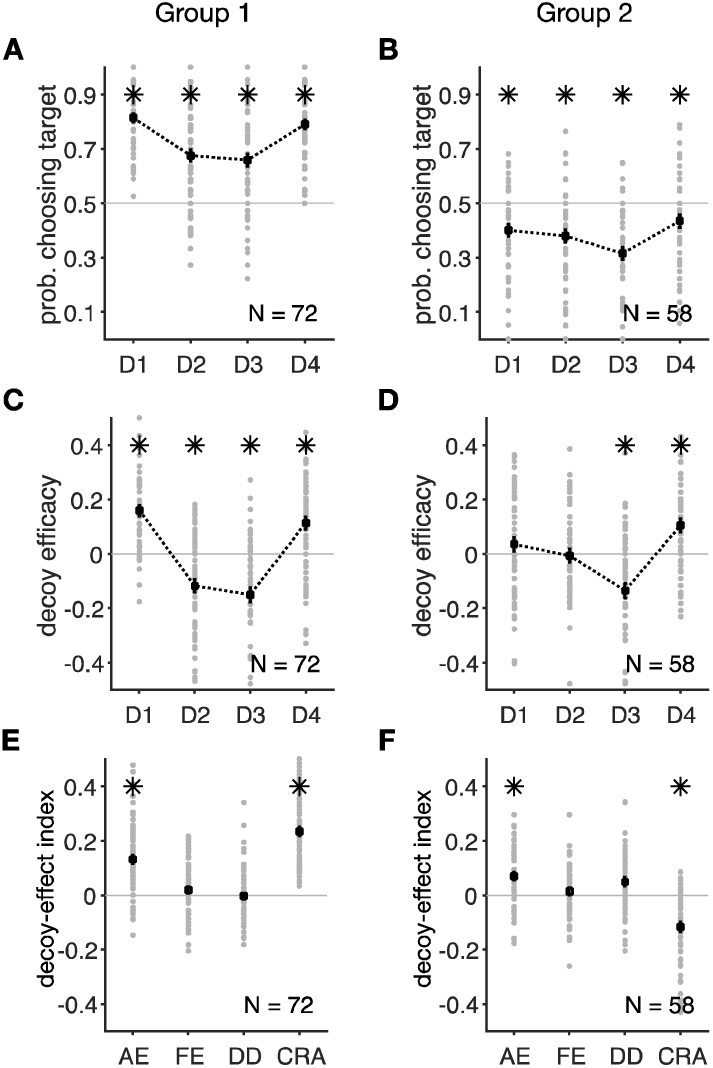
Distinct decoy effects in two groups of subjects identified by clustering based on all decoy-effect indices. (**A–B**) Probability of selecting the target for different decoy locations in the two groups of subjects identified by clustering. Conventions are the same as in **Figure 2**. Subjects in the first group (*N* = 72) selected the target with a probability larger than 0.5 for all decoy locations (increased risk aversion), whereas subjects in the second group (*N* = 58) selected the target with a probability smaller than 0.5 (decreased risk aversion). An asterisk shows that the median of choice probability across subjects for a given decoy location is significantly different from 0.5 (two-sided Wilcoxon signed-test, *p* < 0.05). (**C–D**) Decoy efficacies in the two groups of subjects. Subjects in Group 1 exhibited strong, consistent decoy effects (C), whereas the decoy effects were inconsistent and limited to decoys next to the less risky gamble (target) in Group 2 (D). (**E–F**) Plot shows decoy-effect indices for individuals in the two groups of subjects. The first group showed a strong attraction effect and an overall increase in risk aversion. The second group showed a significant attraction effect and a decrease in risk aversion.

Using ANOVA for each group of subjects, we found significant effects of the decoy location on the average target probabilities (one-way ANOVA; Group 1: *F*(3,284) = 16.8, *p* = 4.7 × 10^−10^; Group 2: *F*(3,228) = 4.95, *p* = 0.002) and decoy efficacies (one-way ANOVA; Group 1: *F*(3,284) = 48.8, *p* = 1.9 × 10^−25^; Group 2: *F*(3,228) = 16.1, *p* = 1.6 × 10^−6^;) in both groups. In addition, Tukey’s *post-hoc* tests revealed significant differences between all decoy location pairs in terms of the average target probabilities and decoy efficacies in both groups (**Supplementary Table 2**).

Finally, both groups showed positive attraction effects (two-sided Wilcoxon signed-test; Group 1: *p* = 0.026, *d* = 1.00; Group 2: *p* = 0.031, *d* = 0.62), but there were no significant frequency effects (two-sided Wilcoxon signed-test; Group 1: *p* = 0.71, *d* = 0.19; Group 2: *p* = 0.51, *d* = 0.15) or differences between dominant and dominated decoys (two-sided Wilcoxon signed-test; Group 1: *p* = 0.64, *d* = −0.04; Group 2; *p* = 0.33, *d* = 0.41). Importantly, subjects in Group 1 exhibited a significant positive change in risk aversion (CRA; two-sided Wilcoxon signed-test, *p* = 0.011, *d* = 1.77; **Figure 4E**), whereas this change was significantly negative for Group 2 (twosided Wilcoxon signed-test, *p* = 0.042, *d* = −0.82; **Figure 4F**). In other words, Group 1 became more risk aversive due to decoy presentation while Group 2 became more risk seeking. However, considering the number of comparisons (N = 4) for each group, only the CRA in Group 1 was statistically significant.

To further reveal the differences in decoy effects between the two groups, we directly compared the four decoy-effect indices between the two groups of subjects. We found a larger overall attraction effect in Group 1 (two-sided Wilcoxon ranksum test, *p* = 0.021, *d* = 0.50; **Figure 5A**), reflecting more consistent decoy effects in this group of subjects. In contrast, the DD was slightly but significantly larger in Group 2 (two-sided Wilcoxon ranksum test, *p* = 0.037, *d* = - 0.48), but this effect was not significant in either group (**Figure 4E-F**). Also, there was no difference between the two groups in terms of the overall frequency effect (two-sided Wilcoxon ranksum test, *p* = 0.49, *d* = 0.05). Finally, the change in risk aversion was much larger in Group 1 than Group 2 (two-sided Wilcoxon ranksum test, *p* = 0.0007, *d* = 2.55).

**Figure 5.**
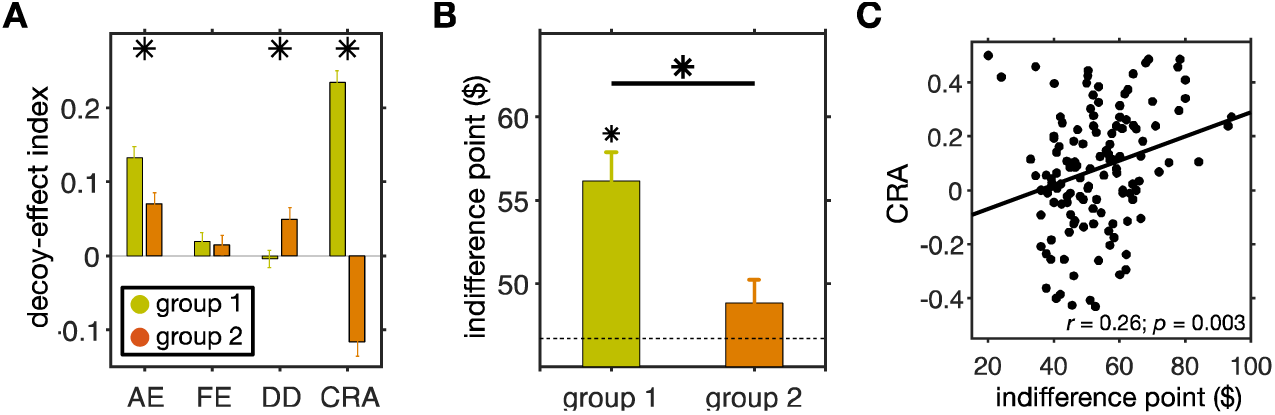
Comparison of the average decoy effects and original risk aversion between the two groups of subjects identified by clustering based on all four decoy-effect indices. (**A**) Plot shows the mean (±std) of each measure separately for the two groups of subjects. An asterisk indicates that the difference between a given measure for the two groups is significant (two-sided Wilcoxon ranksum test, *p* < 0.05). The two groups were different in terms of the overall attraction effect and the overall change in risk preference. (**B**) Comparison between the indifference points for the two groups of subjects. Plot shows the mean (±std) of the indifference point separately for the two groups of subjects and the dashed line indicates an indifference point of $46.70, corresponding to risk-neutrality. The asterisk above the bar shows a significant difference from $46.70 (two-sided Wilcoxon signed-rank test, *p* < 0.05), and the asterisk above the horizontal line indicates a significant difference between the two groups (two-sided Wilcoxon ranksum test, *p* < 0.05). (**C**) Plotted is the change in risk aversion in the decoy task as a function of the indifference point in the estimation task. There was a significant correlation between these quantities indicating that individuals’ risk aversion in the binary choice influences their response to decoys.

Finally, we also examined possible differences in subjects’ behavior under different experimental conditions. To that end, we performed two additional analyses to examine possible differences in behavioral phenotypes between the three cohorts of subjects: control only (N = 38), range-normalization only (N = 48), and mixed cohorts (N = 22). First, we performed clustering analyses for each cohort of subjects separately. For this analysis, we combined data from subjects who performed the control condition first with those who performed the control condition only (combined-control data). We also combined data from subjects who performed the range-normalization condition first with those who performed the range-normalization only (combined-RN data). Comparing average behavior across the two groups of subjects did not reveal any evidence for a difference between patterns of decoy effects in combined-control and combined-RN datasets (**Supplementary Figures 3** and **4**; comparing decoy-effect indices using two-sample t-test, Group 1 AE: *p* = 0.43; Group 1 FE: *p* = 0.19; Group 1 DD: *p* = 0.22; Group 1 CRA: *p* = 0.44; Group 2 AE: *p* = 0.62; Group 2 FE: *p* = 0.49; Group 2 DD: *p* = 0.37; Group 2 CRA: *p* = 0.27). The observed pattern of decoy effects for these datasets was also similar to those found based on whole-data clustering (compare **Figure 4** with **Supplementary Figures 3** and **4**).

Second, we used the two groups of subjects identified based on whole-data clustering and examined the proportion of subjects from each of the three experimental cohorts in these two groups. Again, we did not find any significant differences between the proportion of the two behavioral phenotypes in the three cohorts of subjects (*χ*^2^ = 0.065; *p* = 0.97; control-only cohort: 16 subjects in Group 1 vs. 22 in Group 2; RN-only cohort: 21 subjects in Group 1 vs. 27 in Group 2; mixed cohort: 10 subjects in Group 1 vs. 12 in Group 2). Overall, these analyses did not provide any credible evidence for a difference between behavioral phenotypes of the three cohorts of subjects.

### Relationship between decoy effects and risk preference

The observed difference between the CRA in the two groups of subjects suggests a relationship between risk preference and change in preference due to decoy presentation. To test this relationship directly, we examined the indifference points from the estimation task (note that indifference points from the estimation task were not used for clustering of decoy effects) as the original risk aversion in the two groups of subjects identified in the decoy task. Interestingly, we found that indifference points of subjects in Group 1 were significantly larger than what was predicted by expected value ($46.70), and thus, these subjects were risk averse (two-sided Wilcoxon signed-rank test; *p* = 3.2 × 10^−6^, *d* = 0.65; **Figure 5B**). In contrast, there was no evidence to support that the indifference points of subjects in Group 2 were different from risk neutral (two-sided Wilcoxon signed-rank test; *p* = 0.17, *d* = 0.23). In addition, the indifference points for Group 1 were indifference points of subjects in Group 2 were different from risk neutral (two-sided Wilcoxon significantly larger than Group 2 (two-sided Wilcoxon ranksum test, *p* = 0.0013, *d* = 0.59). These results were completely unexpected because we had not considered indifference points from the estimation task for our clustering.

Considering that the most contrasting differences between the two groups were the change in risk aversion in the decoy task and the indifference points in the estimation task, we examined the relationship between these two quantities within individual subjects. Interestingly, we found a significant positive correlation between the CRA and the indifference point (Pearson correlation; *r* = 0.26, *p* = 0.003; **Figure 5C**), such that subjects with larger indifference points exhibited larger changes in risk aversion due to the decoy presentation. This result indicates that subjects who were more risk averse during binary choice became even more risk averse when considering three gambles. We also computed the correlation between individuals’ degree of risk aversion (measured by indifference points) and the CRA within each group of subjects but did not find any statistically significant relationship in either group (Pearson correlation; Group 1: *r* = 0.10, *p* = 0.42; Group 2: *r* = 0.01, *p* = 0.97).

Finally, we performed additional analyses to test for possible differences between the two groups of subjects and the relationship between individuals’ attraction effect and CRA within each group. Comparing stochasticity in choice (measured by *σ*) between the two groups did not reveal any significant difference (Group 1 median =0.93; Group 2 median =0.75; Wilcoxon ranksum test, *p* = 0.7). There was no significant difference between the median reaction times of the two groups of subjects either (Group 1 median = 0.70 sec; Group 2 median = 0.73 sec; Wilcoxon ranksum test, *p* = 0.3). We also tested the correlation between the AE and CRA in each group correlation; Group 1: *r* = −0.31, *p* = 0.055; Group 2: *r* = −0.014, *p* = 0.9). Overall, these but did not find any evidence for a relationship between these quantities in either group (Pearson correlation; Group 1: *r* = −0.31, p = 0.055; Group 2: *r* = −0.014, p = 09). Overall, these results suggest a distinct rather than a continuous pattern of responses to decoy presentation in the two groups of subjects.

### Effects of decoy distance

Because decoy attributes were determined relative to the attributes of the target or competitor adjacent to them, decoys would be farther from the competitor in terms of reward magnitude for more risk-averse subjects (those with larger indifference points). Considering previous observations that decoy effects increase with the distance of decoy (Soltani et al., 2012), it is possible that more consistent decoy effects in Group 1 subjects was due to larger indifference points for these subjects. We performed additional analyses to examine this possible confound and to test the effect of distance on decoy effects.

First, we divided subjects in each group into two subgroups based on their indifference points (using median) and computed decoy efficacies for each subgroup. However, we did not find any significant difference between decoy efficacies of the two subgroups of either Group 1 or Group 2 (two-sided Wilcoxon ranksum test, *p* > 0.05; **Figure 6A–B**). Therefore, we did not find any evidence that the indifference point influenced decoy effects within each group of subjects.

**Figure 6.**
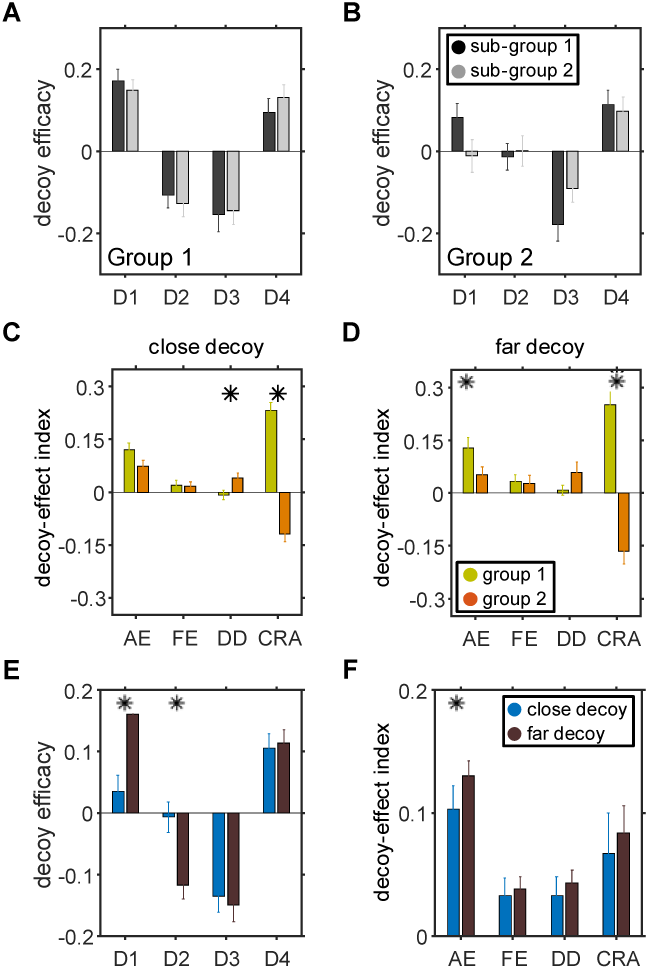
Distinct decoy effects in two groups of subjects identified by clustering does not depend on indifference points or distance of decoys. (**A–B**) Comparisons between the decoy efficacies in subgroups of subjects with small (sub-group 1) or large (sub-group 2) indifference points. Overall, there were no significant differences between decoy efficacies in the two subgroups of either Group 1 or Group 2 (two-sided Wilcoxon ranksum test, *p* > 0.05). (**C–D**) Comparison of the decoy-effect indices between the two groups of subjects identified by clustering, separately for close (C) and far (D) decoys. Plots show the mean (±std) of each measure separately for the two groups of subjects. An asterisk indicates that the difference between a given measure for the two groups is significant (twosided Wilcoxon ranksum test, *p* < 0.05). (**E–F**) Comparison of decoy effects for close and far decoys across all subjects. Plots show the mean (±std) of decoy efficacy (E) and decoy-effect indices (F), separately for close and far decoys. An asterisk indicates that the difference between a given measure for the two conditions is significant (two-sided Wilcoxon ranksum test, *p* < 0.05).

Second, we measured the effect of distance on decoy effects by dividing decoys into far and close decoys (median split based on the absolute Euclidean distance between decoy and the gamble next to it) and calculating decoy-effect indices in each group. We found similar differences between the two groups for close and far decoys (compare **Figure 6C–D** and **Figure 5A**). More specifically, the two groups were statistically different in terms of the overall change in risk preference for both close and far decoys (two-sided Wilcoxon ranksum test; CRA close decoy: *p* = 1 × 10^−21^, *d* = 1.95; CRA far decoy: *p* = 2 × 10^−12^, *d* = 1.35), differential decoy effect for close decoys (two-sided Wilcoxon ranksum test; DD close decoy: *p* = 0.007, *d* = −0.43; DD far decoy: *p* = 0.09, *d* = −0.28), and the overall attraction effect for far decoys (two-sided Wilcoxon ranksum test; AE close decoy: *p* = 0.1, *d* = 0.34) AE far decoy: *p* = 0.04, *d* = 0.36). There was no significant difference between the frequency effect of Groups 1 and 2 for the close or far decoys (two-sided Wilcoxon ranksum test; FE close decoy: *p* = 0.9, *d* = 0.03; FE far decoy: *p* = 0.4, *d* = 0.03).

Finally, we also examined an overall effect of distance by computing decoy efficacies and decoy-effect indices for close and far decoys across all subjects. We found significantly stronger decoy efficacy for far compared with close decoys when they were presented at D1 and D2 locations but not D3 and D4 (two-sided Wilcoxon ranksum test; D1: *p* = 0.0005, *d* = −0.69, D2: *p* = 0.003, *d* = −0.59; D3: *p* = 0.9, *d* = −0.07; D4: *p* = 0.7, *d* = −0.04; **Figure 6E**). Therefore, we found a more consistent distance effect for decoys next to the competitor gamble. In addition, we found the overall attraction effect to be significantly larger for far compared with close decoys (two-sided Wilcoxon ranksum test; AE: *p* = 0.004, *d* = −0.15, FE: *p* = 0.9, *d* = −0.04, DD: *p* = 0.7, *d* = −0.07, CRA, *p* = 0.3, *d* = −0.05; **Figure 6F**). Together, these results suggest that observed differences between the two groups identified by clustering are not likely caused by the differences in the distance of decoys from the competitor.

### New computational model to reveal the underlying mechanisms

In order to explain the observed pattern of choice behavior, we extended the previous range-normalization model of context-dependent choice by Soltani et al. (2012) to include the influence of high-level processes and used this model to fit choice behavior of subjects during the decoy task. The model incorporated both low-level and high-level processes (**Supplementary Figure 5**). Low-level processes correspond to adjustments in reward attribute representations (reward magnitude and probability) by the set of options presented on each trial. We assume linear encoding with threshold and saturation points and implemented low-level processes by dynamically adjusting the slope, threshold, and saturation points based on representation factors and the set of presented gambles on each trial. High-level processes were assumed to operate on the output of neurons encoding gamble attributes in order to change the contribution of each attribute to the final choice. These processes were implemented by allowing different weights for each attribute (gain modulation) based on the configuration of the three gambles in the attribute space, which is assumed to be mediated by attention. Finally, we considered possible combinations for how low- and high-level processes could be implemented with different numbers of parameters (e.g., similar or different representation factors for the two reward attributes), resulting in 17 different models.

We compared these 17 models, which differ in terms of how representations and competitive weighting are modulated, with regards to their ability to account for the pattern of decoy-effect indices in individual subjects in order to determine the best model (see Materials and Methods for more details). We then used the best model parameters to examine relationships between risk preference, adjustments in representations of reward attributes (probability and magnitude), and competitive weighting of these two attributes.

Using three measures for goodness-of-fit, cross-validation prediction error, Akaike information criterion (AIC), and Bayesian information criterion (BIC), we found that a model with both low-level and high-level components provided the best fit to individual subjects’ choice data (**Table 1**). Although overall small values for the cross-validation prediction error could give the wrong impression that different models provide equally good fits, small differences in this measure correspond to large differences in the ability of models to capture choice behavior. Specifically, we found that both AIC and BIC were significantly different between the best model with low-level components only and the best model with both low-level and high-level components (**Table 1**). Therefore, models with and without high-level mechanisms exhibited large differences in terms of their ability to capture choice behavior. The best model (Model # 4 in **Table 1**) required similar location-dependent biases (high-level component) for decoys next to the target and competitor and separate representation factors (*f*_p_ and *f*_m_). The fit based on this model also closely captured the observed pattern of decoy effects in the two groups of subjects (**Figure 7**).

**Table 1.** Comparison of different models’ abilities in capturing subjects’ patterns of decoy efficacies based on three goodness-of-fit measures. Reported are cross-validation prediction error (i.e., the absolute difference between the predicted and actual), AIC, and BIC values for each model and its corresponding sets of parameters. The green shading indicates the best overall model, and blue shading shows the best model with only low-level components. As parameters, σ measures the stochasticity in choice, *f*_0_ is the single neural representation factor, and *f*_*p*_ and *f*_*m*_ are independent neural representation factors for probability and magnitude, respectively. *f*_*pij*_ (respectively, *f*_*mij*_) indicates location-dependent representation factors for probability (respectively, magnitude) with similar values for decoys at locations i and j. High-level parameters *b*_0_ and *b*_*k*_ (*k* = {1, 2, 3, 4}) indicate the constant and location-dependent biases, respectively, and determine the weights of different attributes on final choice (*b*_*ij*_ indicates the case in which location-dependent biases *b*_*i*_ and *b*_*j*_ are assumed to have the same value, *b*_*i*_ = *b*_*j*_ = *b*_*ij*_).

| Model # | Number of parameters | Model parameters | Cross-validation prediction error | AIC | BIC |
| --- | --- | --- | --- | --- | --- |
| 1 | 7 | $f_p, f_m, b_1, b_2, b_3, b_4, \sigma$ | 0.3716 | 285.84 | 305.91 |
| 2 | 5 | $b_1, b_2, b_3, b_4, \sigma$ | 0.0456 | 211.77 | 226.10 |
| 3 | 7 | $f_{p12}, f_{m12}, f_{p34}, f_{m34}, b_{12}, b_{34}, \sigma$ | 0.0371 | 128.28 | 148.35 |
| 4 | 5 | $f_p, f_m, b_{12}, b_{34}, \sigma$ | 0.0344 | 127.53 | 141.86 |
| 5 | 5 | $f_p, f_m, b_{13}, b_{24}, \sigma$ | 0.0433 | 205.53 | 219.86 |
| 6 | 5 | $f_p, f_m, b_{14}, b_{23}, \sigma$ | 0.0476 | 223.96 | 238.29 |
| 7 | 6 | $f_p, f_m, b_{12}, b_{34}, b_0, \sigma$ | 0.0359 | 128.06 | 145.26 |
| 8 | 6 | $f_p, f_m, b_{13}, b_{24}, b_0, \sigma$ | 0.0472 | 223.12 | 240.32 |
| 9 | 6 | $f_p, f_m, b_{14}, b_{23}, b_0, \sigma$ | 0.0493 | 230.05 | 247.25 |
| 10 | 5 | $b_0, f_0, b_{12}, b_{34}, \sigma$ | 0.0521 | 231.08 | 245.41 |
| 11 | 4 | $f_0, b_{12}, b_{34}, \sigma$ | 0.0538 | 237.63 | 249.10 |
| 12 | 4 | $b_0, f_p, f_m, \sigma$ | 0.4007 | 303.31 | 314.78 |
| 13 | 4 | $b_0, b_{12}, b_{34}, \sigma$ | 0.0402 | 181.30 | 192.77 |
| 14 | 3 | $f_p, f_m, \sigma$ | 0.0382 | 169.93 | 178.53 |
| 15 | 3 | $b_{12}, b_{34}, \sigma$ | 0.0459 | 214.97 | 223.57 |
| 16 | 3 | $b_0, f_0, \sigma$ | 0.0502 | 227.08 | 235.68 |
| 17 | 2 | $b_0, \sigma$ | 0.0461 | 213.70 | 219.43 |

**Figure 7.**
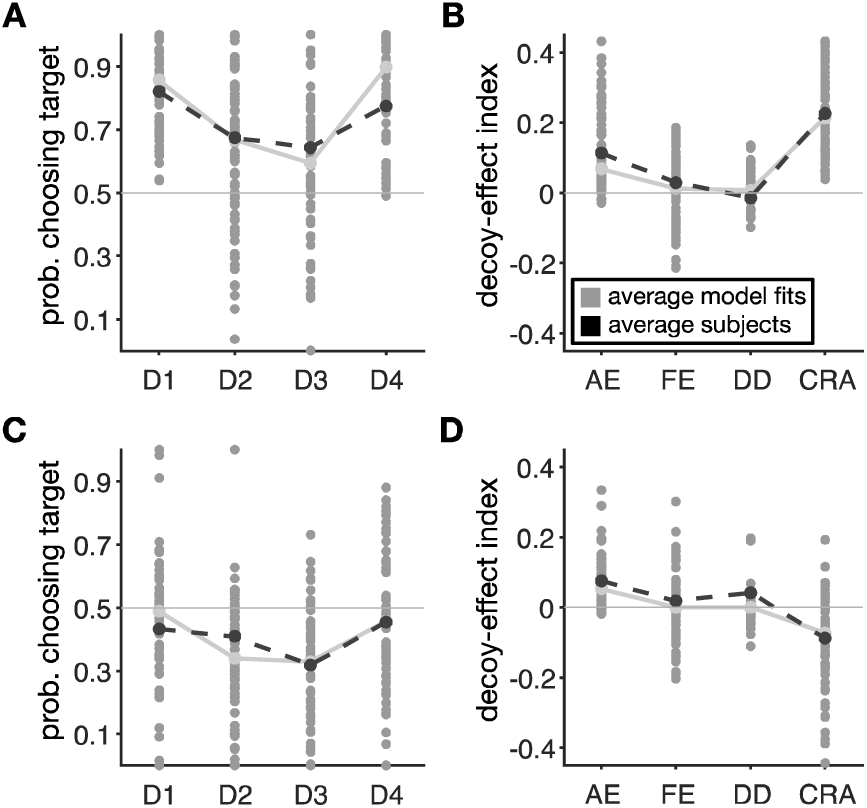
The model with both adjustments of value representation and competitive weighting of reward attributes can capture the experimental data. (**A**, **C**) Each gray circle shows the predicted probability of selecting the target for different decoy locations for individual subjects in the first (A) and second (C) groups using the best overall model. For comparison, the average values across subjects and their predicted values are plotted as well. (**B**, **D**) Plots show predicted decoy-effect indices for individual subjects in the first (B) and second (D) groups of subjects using the best overall model (AE: attraction effect; FE: frequency effect; DD: dominant vs. dominated; and CRA: change in risk aversion).

Despite providing clear results on the best model that can account for experimental data, model selection relies on evaluating the ability of candidate models to predict this data. Other important methods for evaluating computational models are to test their ability to generate the observed behavioral effect and model falsification, which are often ignored (Palminteri et al., 2017). As mentioned above, the best model uses both low-level and high-level processes and is able to replicate the main experimental results (**Figure 7**). To further illustrate the importance of both processes, we simulated behavior in the decoy task using models with only low-level adjustments of neural representations (**Supplementary Figure 6**) or high-level adjustments (**Supplementary Figure 7**) and found that both models failed to capture all aspects of our data. We note that small differences between fits presented in **Figure 7** and **Supplementary Figures 6 and 7** could be misleading because these figures show the average fit of choice data across all subjects and not the individual subjects for which the fit was done. Finally, we also performed model recovery by fitting simulated data and examining estimated model parameters (see *Model recovery and falsification* in Materials and Methods). Overall, we found that our fitting method is able to identify the best model among a set of alternative models (**Supplementary Figures 8**) and moreover, provides an unbiased estimate of model parameters (**Supplementary Figure 9**).

Considering that the best model included both low-level and high-level neural processes, we then investigated the individual effects of adjustments of neural representations and competitive weighting of reward attributes in order to explain the observed pattern of CRA in each group of subjects. Specifically, changes in risk preference due to decoy presentation (measured by CRA) could happen due to different but non-exclusive factors: changes in representations of reward probability and magnitude or changes in how reward probability and magnitude are combined. We compared different models’ behavior and examined estimated model parameters to determine which of the two main mechanisms (change in representations or competitive weighting of reward attributes) contributed to the observed changes in behavior more strongly.

**Figure 8.**
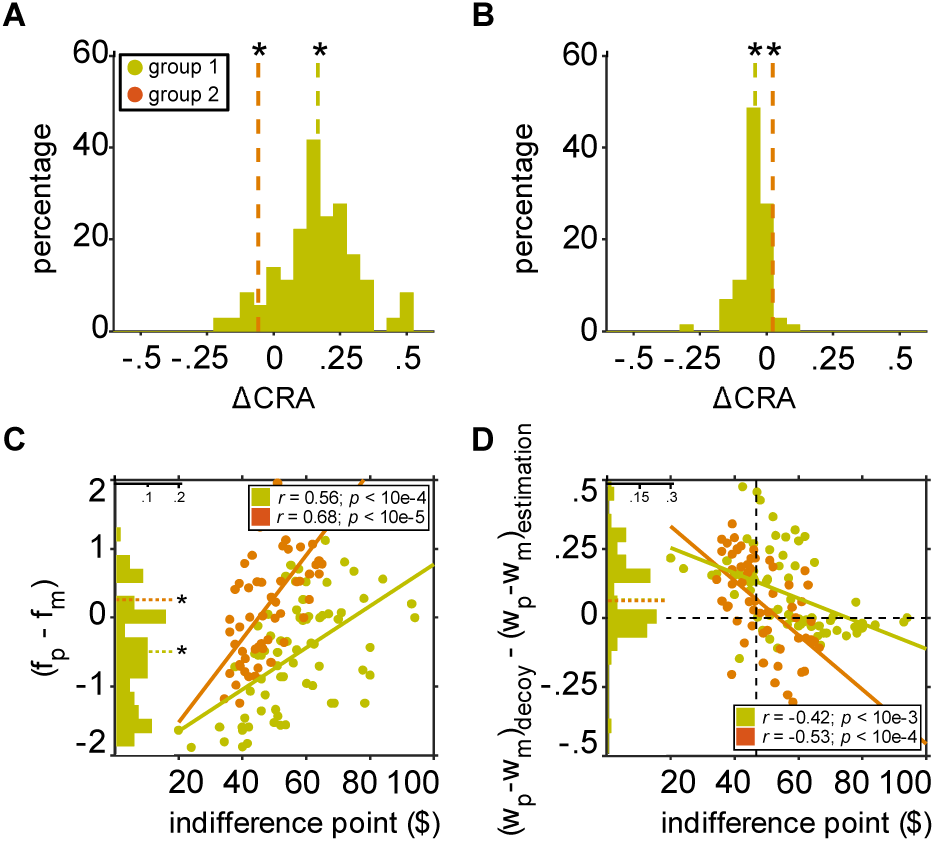
Adjustments of neural representations to decoy presentation and competitive weighting of reward attributes account for opposite patterns of change in risk preference. (**A**) Plotted are the distributions of the difference in CRA between the models with and without adjustments of neural representations (i.e. explained CRA due to neural adjustments) separately for each group of subjects. The dashed lines show the medians, and an asterisk shows a significant difference from zero (twosided Wilcoxon signed-rank test, *p* < 0.05) (**B**) Plotted are the distributions of the difference in CRA between the models with and without competitive weighting of reward attributes (i.e. explained CRA due to competitive weighting) separately for each group of subjects. Both these models include adjustments of neural representations. (**C**) Plotted is the difference of estimated representation factor for probability and magnitude as a function of the indifference point. The green and gray orange histograms plot the fractions of subjects (green: Group 1; orange: Group 2) with certain values of Δf. (**D**) Plotted is the change in the differential weighting of reward probability and magnitude between the estimation and decoy tasks as a function of the indifference point. Adjustments of neural representations to decoy presentation and competitive weighting of reward attributes were both strongly correlated with the original degree of risk aversion. The vertical sashed line indicates the indifference point of $46.70, corresponding to risk-neutrality.

First, we calculated the difference in CRA between the models with and without adjustments of neural representations to decoy presentation. We found that for subjects in Group 1, the change in CRA due to inclusion of low-level neural adjustments was significantly larger than zero (twosided Wilcoxon signed-rank test; median = 0.17, *p* = 6.6 × 10^−12^; **Figure 8A**). In contrast, for subjects in Group 2, the change in CRA due to inclusion of low-level neural adjustments was significantly smaller than zero (two-sided Wilcoxon signed-rank test; median = −0.06, *p* = 0.047). These results are consistent with the observed pattern of CRA in the two groups, indicating that adjustments of neural representations to decoy presentation can account for most of the observed changes in risk preference (increase and decrease in risk aversion in Groups 1 and 2, respectively).

We also examined the relationship between the estimated representation factors (*f*_*p*_ and *f*_*m*_) based on the best model fit and individuals’ degree of risk aversion (indifference points). The representation factor for a given reward attribute determines the dynamic range for coding that attribute or how that reward attribute is represented and adjusted to decoy presentation. More specifically, positive values of the representation factor for a given attribute allow the dynamic range to extend beyond the minimum and maximum of that attribute’s values, whereas the negative values limit the dynamic range between the minimum and maximum values. We found opposite relationships between indifference points and estimated representation factors for probability and magnitude, reflecting competitive adjustments in how the two reward attributes were processed in the decoy task (**Supplementary Figure 10**). Importantly, we found an overall positive correlation between the difference in representation factors (Δf = *f*_p_ − *f*_m_) and the indifference points in both groups of subjects (Pearson correlation; *r* = 0.56, *p* = 3.5×10−5; *r*= 0.68, *p* =4.0×10-6, for Groups 1 and 2, respectively; **Figure 8C**), and this relationship was stronger for Group 2 (two-sided t-test, *p* = = 7.3 × 10^−6^). We focused on the difference between the two representation factors because this difference determines relative processing of the two attributes and thus, the final choice.

These results indicate that in both groups, more risk-averse subjects (subjects with larger indifference points in the estimation task) exhibited more sensitive processing of reward magnitude than reward probability. These results dovetail with the correlation between the CRA and the indifference points (**Figure 5C**), indicating that the adjustment of neural representations was the main factor for driving changes in risk aversion due to decoy presentation. In addition, we found that the differences in the representation factors (Δf) were overall positive in Group 2 (two-sided Wilcoxon signrank test, *p* = 0.03; **Figure 8C** inset) and negative in Group 1 (twosided Wilcoxon signrank test, *p* = 0.008). Note that a larger value for Δf translates to shallower representation for reward probability than reward magnitude, and thus, our results can explain the overall risk attitude in the two groups of subjects.

Second, we calculated the difference in CRA between the models with and without competitive weighting of reward attributes in order to determine how much of the CRA was influenced by this high-level process (both these models include adjustments of neural representations). We found that for subjects in Group 1, the overall change in CRA due to inclusion of competitive weighting was significantly smaller than zero (two-sided Wilcoxon signed-rank test; median = - 0.04, *p* = 4.8 × 10^−7^; **Figure 8B**). In contrast, for subjects in Group 2, the change in CRA due inclusion of competitive weighting was significantly greater than zero (two-sided Wilcoxon signed-rank test; median = 0.02, *p* = 2.0 × 10^−6^). These results indicate that competitive to weighting gives rise to a decrease (increase) in risk aversion in Group 1 (respectively, Group 2), which is the opposite of the effect of adjustments in neural representations. Therefore, competitive weighting of reward attributes, which closely resembles selective attentional modulation, can partially compensate for the divergence of risk preference in the two groups of subjects (**Figure 8A**) due to adjustments in value representation.

Interestingly, competitive weighting of reward attributes was also strongly correlated with the original degree of risk aversion (indifferent point). Specifically, we found significant negative correlations between the changes in the differential weighting of reward probability and magnitude between the estimation and decoy task ([*w*_*p*_ – *w*_*m*_]_*decoy*_-[*w*_*p*_ –*w*_*m*_]_*estimation*_) and the indifference points (Pearson correlation; *r* = −0.42, *p* = 8.2 × 10^−4^; *r* = −0.53, *p* = 7.8 × 10^−5^, for Groups 1 and 2, respectively; **Figure 8D**). This relationship was stronger for Group 2 (two-sided t-test, *p* = 0.003) indicating that similar to low-level neural adjustments, high-level changes in attribute weighting were also more strongly correlated with the original risk aversion in this group. Note that *w*_*p*_ and *w*_*m*_ measure the weights of reward probability and magnitude on the final choice, respectively, and thus the differential weighting of reward probability and magnitude ([*w*_*p*_ −*w*_*m*_]_*estimation*_) measures the degree of risk aversion in the binary gambling (estimation) task for each individual. As a result, a decrease in the differential weighting of reward probability and magnitude between the estimation and decoy tasks ([*w*_*p*_ − *wm]decoy*− *[wp*−*wm]estimation)<0)* would give rise to more risk-seeking behavior and vice versa. The negative correlation between changes in differential weighting of reward probability and magnitude and the original degree of risk aversion shows that high-level adjustments to decoy presentation (due to competitive weighting of reward attributes) can push risk preference toward risk neutrality.

Together, results of fitting based on our model reveal how two separate mechanisms drive subjects’ risk behavior during multi-attribute choice and give rise to opposite changes in risk preference. On the one hand, low-level adjustments of value representation cause more risk aversion proportional to individuals’ degree of risk aversion in the binary gambling task. On the other hand, high-level competitive weighting of reward attributes gives rise to more risk-seeking behavior by increasing the relative weight for probability and magnitude inversely proportional to individuals’ original degree of risk aversion.

## Discussion

Previous studies on choice between multiple options have illustrated that context effects are general features of human choice behavior (Tversky, 1972; Huber et al., 1982; Bettman et al., 1998; Dhar and Simonson, 2003; Simonson 1989; Bettman et al., 1998; Kahneman and Tversky, 1984; Doyle et al., 1999) and moreover, revealed great individual variability across different types of choice (Huber and Puto, 1983; Wedell and Pettibone, 1996; Trueblood, 2012; Soltani et al., 2012; Pan and Lehmann, 1993; Pettibone and Wedell, 2007; Ha et al., 2009), indicating that multiple processes are involved in context-dependent choice. Here, we used choice between multiple risky options to study how context effects are influenced by individuals’ risk attitudes and neural adaptation on different timescales. With a large cohort of subjects and a new computational model, we explored individual variability in how reward information is processed and how it influences context effects.

To our surprise, we did not observe any evidence for the effects of neural adaptation on the order of 100 trials on decoy effects. Although this null result could be due to many factors, previous studies suggest that value representations adapt to the range of reward values available in a given condition (e.g., block of trials). For example, Padoa-Schioppa (Padoa-Schioppa, 2009) has shown that the sensitivity of orbitofrontal cortex (OFC) neurons encoding reward value is inversely proportional to the range of values for the available options, and this adaptation can result in optimal coding of reward value (Rustichini et al., 2017). Moreover, neural adaptation is prevalent in different brain areas (Albright and Stoner, 2002; Tobler et al., 2005; Sallet et al., 2007; Carandini and Heeger, 2012; Summerfield and Lange, 2014) and could strongly influence context effects. Interestingly, two recent studies have shown that OFC neurons signal context-dependent reward value only during free choice (Yamada et al. 2018) and contribute to decision making in different contexts (Xie and Padoa-Schioppa, 2016), indicating that neural adaptation in OFC could contribute to value-based decision making.

The lack of evidence for the contribution of long-term neural adaptation to context effects in our data could be due to many factors including ineffectiveness of our manipulation, individual variability, etc. Nonetheless, our results suggest that certain neural computations, such as long-term adaptations to the range of reward attributes, are performed only when they allow more flexible behavior (Soltani et al., 2016; Farashahi et al., 2018; Soltani and Izquierdo, 2019; Spitmaan et al., 2019). Moreover, context effects could be influenced by adaptations to other aspects of reward information such as mean, variance, and skewness, none of which was manipulated in our study. Despite finding null results for range manipulation, the control and range-manipulation conditions together provided us with a large dataset to investigate individual variability and behavioral phenotype with respect to decoy effects.

Using our large dataset, we found that context effects strongly depend on individuals’ degree of risk aversion in the binary gambling task. Specifically, using a clustering method, we identified two groups of subjects with different patterns of decoy effects. Subjects in the one group exhibited consistently significant decoy effects and increased risk aversion due to decoy presentation. In contrast, subjects in the other group did not show consistent decoy effects and became more risk seeking due to decoy presentation. We found that subjects in the first group were more risk averse when choosing between two gambles. Thus, for the first time, to the best of our knowledge, we were able to identify distinct behavioral “phenotypes” in terms of context effects and the influence of risk attitudes on these effects.

We note that despite the *post-hoc* nature of clustering analyses, our findings based on these analyses are valid because of the size of our datasets and because of the cross-validated and agonistic clustering method we used, ensuring that there was no overfitting and no knowledge of data was used to determine the number of clusters, etc. Important to our finding on the relationship between risk attitudes and context effects is that risk aversion was measured during the binary gambling (estimation) task that was performed before the decoy task and its results were not used for clustering of decoy effects. Nevertheless, future experiments can be used to further validate our results. Together, our results suggest that understanding individual variability could be very informative about high-level cognitive functions and their interactions with low-level neural processes.

Changes in risk preference due to decoy presentation (measured by the CRA in this study) could happen due to changes in representations of reward attributes, changes in how reward probability and magnitudes are combined, or both. We incorporated two types of processes in our models (change in representations and competitive weighting of reward attributes) in order to pinpoint which mechanism(s) underlies the observed change in risk preference. Based on the fit of experimental data, we conclude that both types of changes contribute to the observed shifts. We also estimated how much of the CRA is due to changes in representations or due to competitive weighting of reward attributes and found the former to be more important. Understanding how our findings apply to context effects for choice between non-risky options requires additional experiments, but a similar computational approach can be applied to understand the results. We speculate that factors such as saliency of certain attributes or loss aversion could similarly drive attention when more than two options are available, resulting in a shift in those measured factors when going from binary choice to choice between multiple options.

Although extant models of context-dependent choice assume that context effects occur due to either high-level cognitive or low-level neural processes, we found that both processes are necessary to account for the complex pattern of decoy effects. For example, it has been shown that adjustments of reward value representations can contribute to context effects via different neural mechanisms such as range normalization (Soltani et al., 2012; Louie et al., 2013) or divisive normalization (Louie et al., 2011; Carandini and Heeger, 2012; Heeger, 1992). We found that these adjustments can account for some but not all of the observed context effects. High-level models rely on different cognitive processes to account for context effects, including attentional switching to different choice attributes, menu-dependent evaluation of choice attributes, competition between attribute processing to enhance contrast between certain attributes (Tversky, 1972; Tsetsos et al., 2010; Tversky and Simonson, 1993; Roe et al., 2001; Usher and McClelland, 2004; Johnson and Busemeyer, 2005; Usher et al., 2008; Trueblood et al., 2014; Choplin and Hummel, 2005), and a more recent model based on rank-dependent weighting of information according to the salience of the sampled information (Tsetsos et al., 2012). In addition to adjustments to the set of options on each trial, we found that the model that best fit the experimental data includes competitive weighting of reward probability and magnitude. Therefore, the additional high-level mechanism included in our model shares many general features of previous high-level models. We note that we did not perform model comparison with existing models because our goal here was to link context effects to risk aversion. Therefore, despite being able to reveal this link, our proposed model may not be the only model that can capture our experimental results.

Finally, our model revealed the relationship between context effects and risk preference within individuals. Specifically, we found that both low-level adjustments in value representation due to decoy presentation and high-level adjustments in weighting of reward attributes, perhaps via attentional modulations, were correlated with the original degree of risk aversion within individuals but in the opposite directions. The opposite modulations of low-level and high-level processes shifted risk preference toward risk-aversion and risk-seeking behavior, respectively, and created a distinct pattern of choice behavior in a more complex task.

## Materials and Methods

### Ethics statement

The study was approved by the Institutional Review Board of Dartmouth College. A consent form was obtained from each subject prior to participating the experiment.

### Subjects

In total, 108 healthy Dartmouth College students with normal or corrected to normal vision participated in the study (50 females and 58 males): 38 exclusively in the control condition, 48 exclusively in the range-manipulation condition, and 22 in both conditions, resulting in 130 sets of data. Therefore, 60 and 70 datasets correspond to the control and range-manipulation conditions, respectively. Data from all subjects were included in the data analyses. Subjects were compensated with a combination of money and “t-points,” that are extra-credit points for classes within the department of Psychological and Brain Sciences at Dartmouth College. More specifically, in addition to the base rate of $10/hour or one t-point/hour, subjects were compensated by up to $10/hour depending on the gambles they chose during the tasks (see below).

### Experimental paradigm

Subjects performed two gambling tasks consisting of an estimation and a decoy task. Moreover, there were two conditions within the decoy task: control and range-manipulation. In both tasks, subjects were told to select the gamble they believe would result in more reward considering both options’ reward probabilities and magnitudes. We told the subjects that they would only be paid based on the outcome of one of their randomly selected gambles in each task; therefore, every trial was equally relevant to their final payoff. The estimation task was performed first followed by the control and range-manipulation conditions of the decoy task. The order of the control and range-manipulation conditions was randomized between subjects.

In the estimation task (70 trials), subjects had to choose between two gambles defined by the probability *p* of earning monetary reward *m* (*p, m*) (**Figure 1A**). The two reward attributes green). (probability and magnitude) were represented on the screen using two different colors (red and green). The target (T) gamble was low-risk and had a low reward magnitude (*p*= 0.7, *m* = $20 ± 2, while the competitor gamble (C) was high-risk and had a variable reward magnitude (*p* = 0.3, m = $30 – $80). On each trial of this task, two gambles appeared on the screen with the message “Evaluate” on the top of the screen. Subjects had 4 seconds to evaluate both gambles and decide their preferred gamble. When the 4 seconds were over, the message changed to “Choose,” and subjects had 1 second to make their selection using the keyboard. Once the selection was made, they could see their choice for one second. Subsequently, a fixation cross appeared on the screen indicating that the next trial was about to start. The choice data of the subject in this task was used to determine the subject’s attitudes toward risk. This information was then used to tailor equally preferable target and competitor gambles (for each subject) in the decoy task.

The decoy task was used to determine how the presence of a third gamble changed the preference between the target and competitor gambles. During each trial of the decoy task, three gambles (presented on the same horizontal plane) and a message on top of the screen saying “Evaluate” appeared for 6 seconds (**Figure 1B**). After the evaluation period, the message changed to “Choose” and one of the gambles was simultaneously removed. Subjects had 1 second to choose their preferred gamble from the remaining two using the keyboard. We allowed a short amount of time for submitting a response in order to avoid re-evaluation of the remaining options. After making a choice, their selected gamble appeared for 1 second, followed by a fixation cross indicating that the next trial was going to start. The target and the competitor gambles were tailored for each subject to be equally preferable using the subjective indifference point computed from the estimation task (see below). The third gamble (decoy) could be in one of the following four positions in the attribute space (**Figure 1C**): D1 decoys, which were better than the competitor gamble in terms of both reward probability and magnitude but were better than the target only in terms of magnitude (asymmetrically dominant); D2 decoys, which were worse than the competitor in both probability and magnitude but were worse than the target only in terms of probability (asymmetrically dominated); D3 decoys, which were better than the target in both probability and magnitude but were better than the competitor only in terms of probability (asymmetrically dominant); and D4 decoys, which were worse than the target in both probability and magnitude but were better than the competitor only in terms of probability (asymmetrically dominated). Therefore, decoys could be asymmetrically dominant or asymmetrically dominated. Four decoy types were presented equally (30 times each), and the decoy type was pseudo-randomly assigned in every trial.

Overall, each subject performed 120 trials in the control condition and 150 trials in the range-manipulation condition of the decoy task. In 2/3 of the trials, the decoy gamble disappeared, whereas in the remaining 1/3 of the trials, either the target or the competitor was randomly selected to disappear in order to make decoy identification impractical. Removing the decoy before choice created the so-called phantom decoy effect, as the decoy was not available in the choosing portion of the trial. This design allowed us to measure the effect of asymmetrically dominant decoys that could not be studied otherwise. The trials in which either the target or the competitor disappeared were not considered in the analyses. Because we only allowed only one second for submitting a response, subjects could miss a significant percentage of trials (median of missed trials was equal to 12%). To avoid losing too many trials, we added any missed trials back into the subject’s set such that each subject completed the same number of total trials. However, due to trial replacement, it took longer for subjects (that missed many trials) to perform the experiment, and consequently, some subjects were unable to complete the task. More specifically, we terminated the experiment if it took longer than 60 minutes to complete or if the subject missed more than 15 trials in a row. Overall, these criteria resulted in termination of experiments for 17 unmotivated subjects.

In the control condition, the target was low-risk and had a low reward magnitude (*p* = 0.7 ± 0.05, *m* = $20 ± 2), and the competitor was high-risk and had a variable reward magnitude (*p* = 0.3 ± 0.05,, *m* = $*m*_0_ ± 2), where *m*_0_ is the indifference point (see Eq. 1 below) obtained in the estimation experiment. Specifically, the magnitude value of the competitor was tailored for each subject based on their data from the estimation task such that the competitor and the target were equally preferable for every subject. In addition, reward probability and magnitude of the decoy gambles were determined relative to the locations of adjacent target or competitor gamble in the attribute space. More specifically, we calculated decoy attributes by shifting reward probability and magnitude of the target (or competitor) by ±5%, ±15%, or ±30%. However, to avoid very small and large reward attributes, we constrained reward probabilities and magnitudes to [0.2, 0.85] and [$15, $70], respectively). The range-manipulation condition was similar to the control condition except that 20% of the trials presented a set of “high-range” gambles. The reward probability and magnitude of the high-range gambles were the following: (*p* = 0.9 ± *0*.*05, m=$10 ± 2, p=0*.*1 ± 0*.*05, m=$90 ± 2*, and *(p=0*.*5 ± 0*.*05, m=$20 ± 2)*. The high-range trials were removed from the subsequent analysis, as their only purpose was to expand the range of gamble attributes and disrupt neural adaptation.

### Data analysis

Subjects’ choice behavior in the estimation task was fit using a sigmoid function. More specifically, the probability of choosing the low-risk gamble (the gamble with reward probability *p*_*R*_ = 0.7 and a reward magnitude of *m* = $20), *p*_*low-risk*_, was fit as a function of the reward magnitude of the high-risk gambles, *m*_*high-risk*_:

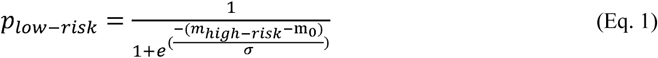

where *m*_0_ corresponds to a reward value at which the high-risk gamble was equally preferable to the low-risk gamble (i.e., the indifference point), and σ measures the stochasticity in choice.

In the decoy task, we computed the decoy efficacy (*DE*_*k*_) for each of four locations in the attribute space that a decoy could appear as follows:

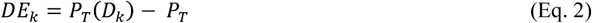

where *P*_*T*_(*D*_*k*_)is the probability of choosing the target given the decoy *k* (*k* = {1,2,3,4}) and *P*_*T*_ location. *P*_*T*_ is calculated as the probability of choosing the target gamble across all four decoy is the probability of choosing the target gamble for a given subject regardless of the decoy locations. Importantly, decoy efficacies allowed us to measure decoy effects for each location relative to the overall subject preference for the target gamble (*P*_*T*_).

In order to fully characterize the change in preference due to the introduction of a decoy to the choice set, we defined four orthogonal quantities (“decoy-effect” indices) based on linear transformations of decoy efficacies and overall target selection as follows:

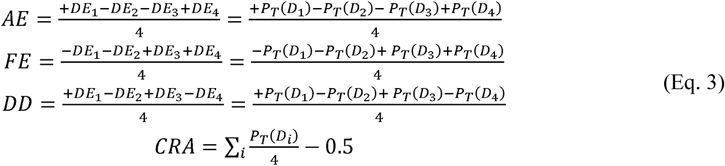

The first index, the overall attraction effect (AE), is computed by averaging decoy efficacies over all decoy locations and considering reversed signs for the asymmetrically dominant and dominated decoys for the target and competitor ([D1-D2] versus [-D3+D4]). On the one hand, presentation of D1 and D4 decoys can increase the preference for the target by asymmetrically dominating the competitor or being dominated by the target, respectively. On the other hand, D2 and D3 decoys can reduce the preference for the target by being asymmetrically dominated by the competitor and dominating the target, respectively. Therefore, the AE measures the overall attraction effect for both dominant and dominated decoys, or more specifically, how much the presentation of the decoy increases preference for the target relative to the competitor. The second index, the overall frequency effect (FE), quantifies the overall tendency to choose the gamble next to the decoy in terms of both reward probability and magnitude (i.e., the competitor for D1 and D2 decoys, and the target for D3 and D4 decoys). The third index, the dominant vs. dominated (DD), quantifies the overall difference between the effects of dominant and dominated decoys. The fourth index measures the overall change in risk aversion (CRA) due to the decoy presentation. Note that due to the symmetry in decoy presentation and because the target and competitor gambles were set based on the indifference point in the estimation task, these gambles should be selected equally (with probability equal to 0.5) in the absence of any changes in risk aversion due to decoy presentation. These four decoy-effect indices are linear transformations of decoy efficacies and could take a value between −0.5 and 0.5. Importantly, these indices are orthogonal to each other.

For analysis of the time course of decoy effects, we used a moving window with a length of 40 trials and a step size of 20 trials in order to calculate changes in decoy-effect indices and the probability of choosing the target over time. The statistical test used for each comparison is reported where it is mentioned, and the reported effect sizes are Cohen’s d values.

### Clustering analyses

To identify distinct patterns of decoy effects, we separated subjects into different clusters using the k-means clustering method based on various sets of decoy-effect indices as features and for different numbers of clusters. Specifically, we tried all possible exclusive combinations of the four decoy-effect indices to cluster subjects. The “silhouette” values were calculated for clustering data with different features and numbers of clusters. The silhouette values range from −1 to 1, where a high value indicates that the data point is well matched to its own cluster and poorly matched to neighboring clusters. We found that clustering based on all four decoy-effect indices and into two clusters provides the best separation between data. We then used the clustering indices based on this model to quantify subjects’ behavior in each cluster/group.

### Computational models

We extended a previous model of context-dependent choice by Soltani et al. (2012) to include the influence of high-level cognitive processes. The previous model only considered changes in representations of reward probability and magnitude to account for decoy effects. In addition to these low-level mechanisms, the new model also considered mechanisms for how reward probability and magnitude are combined and how this combination is modulated by attention, which we assumed to be high-level processes. We assumed that the values of gambles in a given attribute dimension (reward magnitude and probability) are encoded by a neural population selective to that attribute (attribute-encoding neurons). The output of attribute-encoding neurons, in turn, projects to value-encoding neurons that compute the overall value of each gamble (**Supplementary Figure 5**).

More specifically, we assumed that the response of neurons encoding attribute *i* (*i* = {*m,p*}), *r*_*i*_, is a linear function of the attribute value *v*_*i*_ as follows:

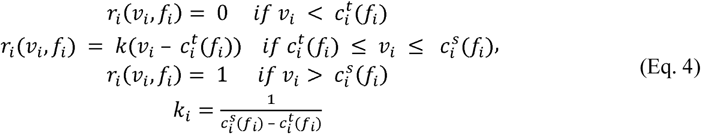

where *k* is the slope, 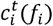 and 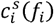 are the threshold and saturation points, respectively and *f*_*i*_ is the representation factor that determines the dynamic range (i.e., the interval between threshold and saturation) of neural response for attribute *i*. Specifically, the threshold and saturation points depend on the representation factor and the attribute values of the gambles in the choice set as follows:

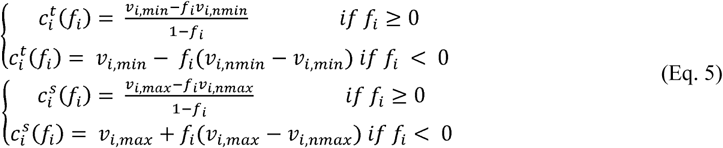

where *v*_*i,min*_ and *v*_*i,nmin*_ are the minimum and next to the minimum values of the presented gamble in a given attribute, respectively; and *v*_*i,max*_ and *v*_*i,nmax*_ are the maximum and the next to the maximum values of the presented gamble, respectively. Positive values of *f*_*i*_ allow the dynamic range to extend beyond the minimum and maximum attribute values, whereas the negative values limit the dynamic range between the minimum and maximum values (Soltani et al., 2012). Therefore, *f*_*i*_’s sign and magnitude determine the dynamic range of neural response.

The overall value of each gamble, *V*(*X*), is computed by a weighted sum of outputs from the attribute-encoding neurons as follows:

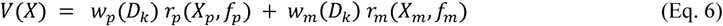

where *w*_*i*_ (*D*_*K*_) is the weight of the connection from the attribute-encoding population *i*(*i* = {*m, p*}) to the option-value population when decoy *D*_*k*_ was presented (*k*={1,2,3,4}).

We considered an additive model instead of the multiplicative one often used in the literature for a few reasons. First, we wanted to make our new model consistent with the model of Soltani et al. (2012). Second, although additive and multiplicative models provide very similar fit of choice behavior, only additive models allow differential weighting of the two reward attributes (e.g., via attentional mechanisms). In a multiplicative model, differential weighting of attributes is only possible by adopting non-linear functions of reward magnitude and probability (i.e., non-linear utility and probability weighting functions) and assuming that these functions––which are assumed to be innate and fixed––change depending on decoy locations. This suggests critical limitations for multiplicative models in explaining flexible choice behavior (Farashahi et al., 2019).

For some model simulations and fitting, we assumed a single neural representation factor *f*_0_ to capture neural adjustments for both reward probability and magnitude:

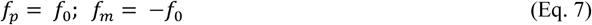

where *f*_*p*_ and *f*_*m*_ are neural representation factors for reward probability and magnitude, respectively. For other simulations and fitting, we used separate representation factors (*f*_*p*_ and *f*_*m*_) to allow more flexible representation.

To determine the weights of the connections between the attribute-encoding and option-value encoding neurons (*w*_*i*_(*D*_*k*_)), we assumed a competitive weighting mechanism as follows:

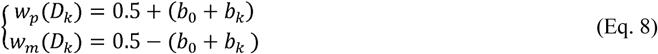

where *b*_0_ is a constant bias and *b*_*k*_ is the location-dependent bias for decoy D_*k*_. The constant bias, *b*_0_, could be a free parameter or could be set based on the choice behavior in the estimation task as follows:

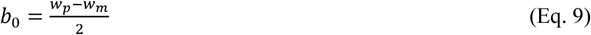

where *w*_*p*_ and *w*_*m*_ are estimated weights for the reward probability and magnitude in the estimation task, respectively (using Eq. 6). Note that the differential weighting of reward probability and magnitude ([*w*_*p*_ – *w*_*m*_]) in the estimation task was strongly correlated with the indifference points (Pearson correlation; *r* = 0.92, *p =* 2.8 × 10^-8^), indicating that the former quantity measures the “original” degree of risk aversion for each subject.

The location-dependent biases, *b*_*k*_, could be different for the four decoy locations or coupled

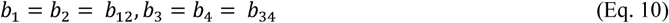

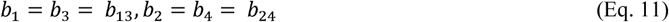

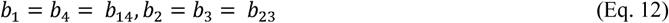

Although we considered all the above possibilities to determine the weights, only coupling based on Eq. 10 does not require identification of decoys and is plausible. Finally, we also assumed separate location-dependent biases for probability and magnitude in order to construct a more-flexible model.

Overall, different choices for setting similar or separate parameters for representation factors and attribute weighting resulted in the generation of 17 models. We used these 17 models to fit individuals’ choice data in terms of decoy-effect indices using three measures for goodness-of-fit. First, we performed cross validation to compare the prediction error of decoy-effect indices to determine the best model. Specifically, we used 70% of data (randomly chosen) to estimate the best-fitting parameters and then calculated the model prediction errors over the remaining 30% of data. One hundred different data partitions were generated for estimation and prediction purposes. We also calculated the Akaike information criterion (AIC) and Bayesian information criterion (BIC) based on all data from each individual subject using the following equations:

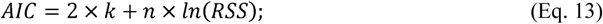

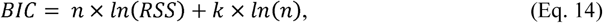

where *k* is the number of parameters for each model, *n* represents the number of total datapoints, and *RSS* indicates the residual sum square of the model. Overall, we found that the model with separate representation factors (*f*_*p*_ and *f*_*m*_) and similar location-dependent biases for decoys next to the target and competitor (Eq. 10) provides the best fit. Therefore, we chose this model to estimate the representation factors and weights of connections for each subject and how they are related to individuals’ risk preference.

### Model recovery and falsification

For model recovery, we generated data for the decoy task using 200 different instances of each model (total 17 models). We then fit the generated data with all models in order to find the best fitting model based on AIC (similar results are obtained using BIC). The results can be summarized by a confusion matrix that plots the percentage of instances that a model used to generate the data was best fit by the same or other models (**Supplementary Figure 8**). We found that the majority of models can predict their own generated data better than the other competing models. We also calculated the estimation error for 5 parameters of the best model (Model # 4 in **Table 1**). For this analysis, we generated 500 sets of model parameters. We did not find any significant difference between the estimation and actual model parameters, indicating that the fitting method can provide an unbiased estimate of model parameters (**Supplementary Figure 9**).

For model falsification and to further show the importance of both processes, we simulated behavior in the decoy task using the best model (**Figure 7**) and models with only low-level adjustments of neural representations (**Supplementary Figure 6**) or high-level adjustments of attribute weights on the final choice (**Supplementary Figure 7**).

## Acknowledgments

We thank Daeyeol Lee and Chanc VanWinkle Orzell for helpful comments on the manuscript.

---

**Supplementary Table 1.** Comparison of the average probability of selecting the target and decoy efficacies between different decoy locations using Tukey’s HSD *post-hoc* test, separately for the control and range-manipulation conditions. Reported are *p*-values for comparison of a pair of decoy locations (rows) and for different quantities and experimental conditions (columns). The orange shading indicates *p*-values that are smaller than 0.05 and thus differences that are statistically significant.

| comparison | prob. choosing target (control) | decoy efficacy (control) | prob. choosing target (range-manipulation) | decoy efficacy (range-manipulation) |
| --- | --- | --- | --- | --- |
| D<sub>1</sub>-D<sub>2</sub> | 0.0767 | 0.0001 | 0.0045 | 0.0001 |
| D<sub>1</sub>-D<sub>3</sub> | 0.0017 | 0.0001 | 0.0230 | 0.0001 |
| D<sub>1</sub>-D<sub>4</sub> | 0.0934 | 0.0495 | 0.0902 | 0.0437 |
| D<sub>2</sub>-D<sub>3</sub> | 0.0142 | 0.0278 | 0.0574 | 0.0301 |
| D<sub>2</sub>-D<sub>4</sub> | 0.0421 | 0.0001 | 0.0124 | 0.0001 |
| D<sub>3</sub>-D<sub>4</sub> | 0.0044 | 0.0001 | 0.0539 | 0.0001 |

**Supplementary Table 2.**
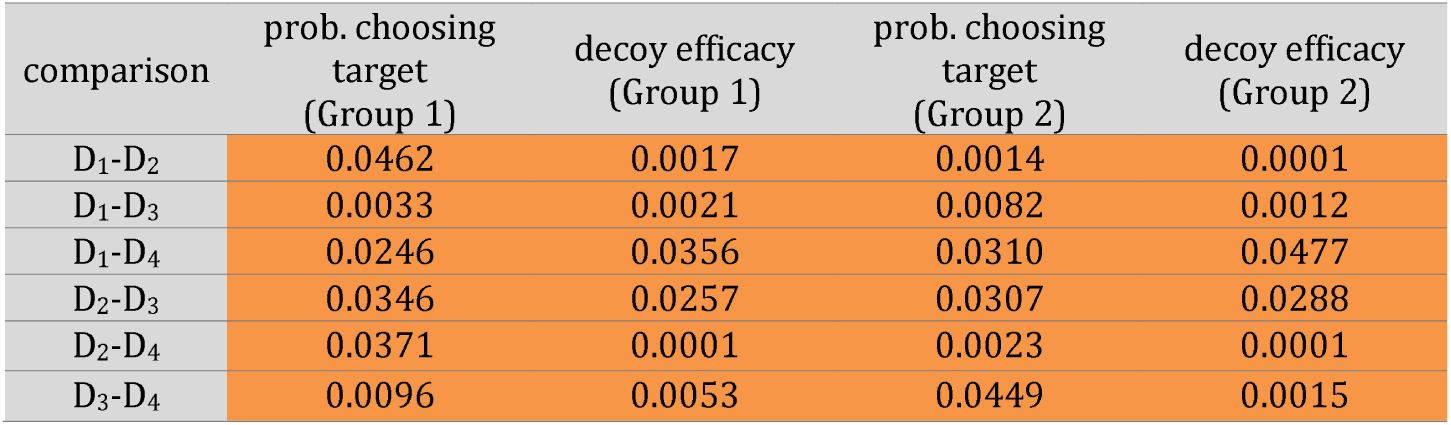
Comparison of the average probability of selecting the target and decoy efficacies between different decoy locations using Tukey’s HSD *post-hoc* test, separately for the two clusters of subjects. Reported are *p*-values for comparison of a pair of decoy locations (rows) and for different groups and quantities (columns). The orange shading indicates *p*-values that are smaller than 0.05 and thus differences that are statistically significant.

**Supplementary Figure 1.**
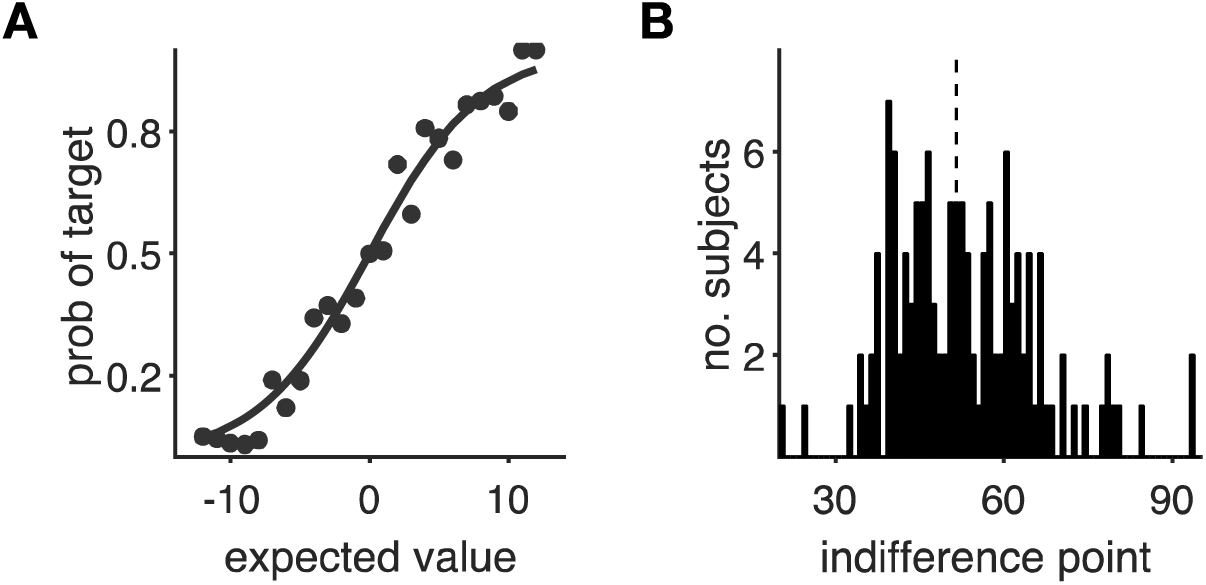
Behavior of subjects during the estimation task. **(A)** Plotted is the probability of choosing the low-risk gamble (target) as a function of the reward magnitude of the high-risk gambles for an individual subject. The solid curve shows the fit using a sigmoid function. Each black dot represents an individual subject’s data. **(B)** Distribution of estimated indifference points in estimation task across subjects. The indifference point is defined as the magnitude of a high-risk gamble that was as equally preferred as the low-risk gamble. The dashed line indicates the median. Overall, we observed large variability for the indifference point across subjects.

**Supplementary Figure 2.**
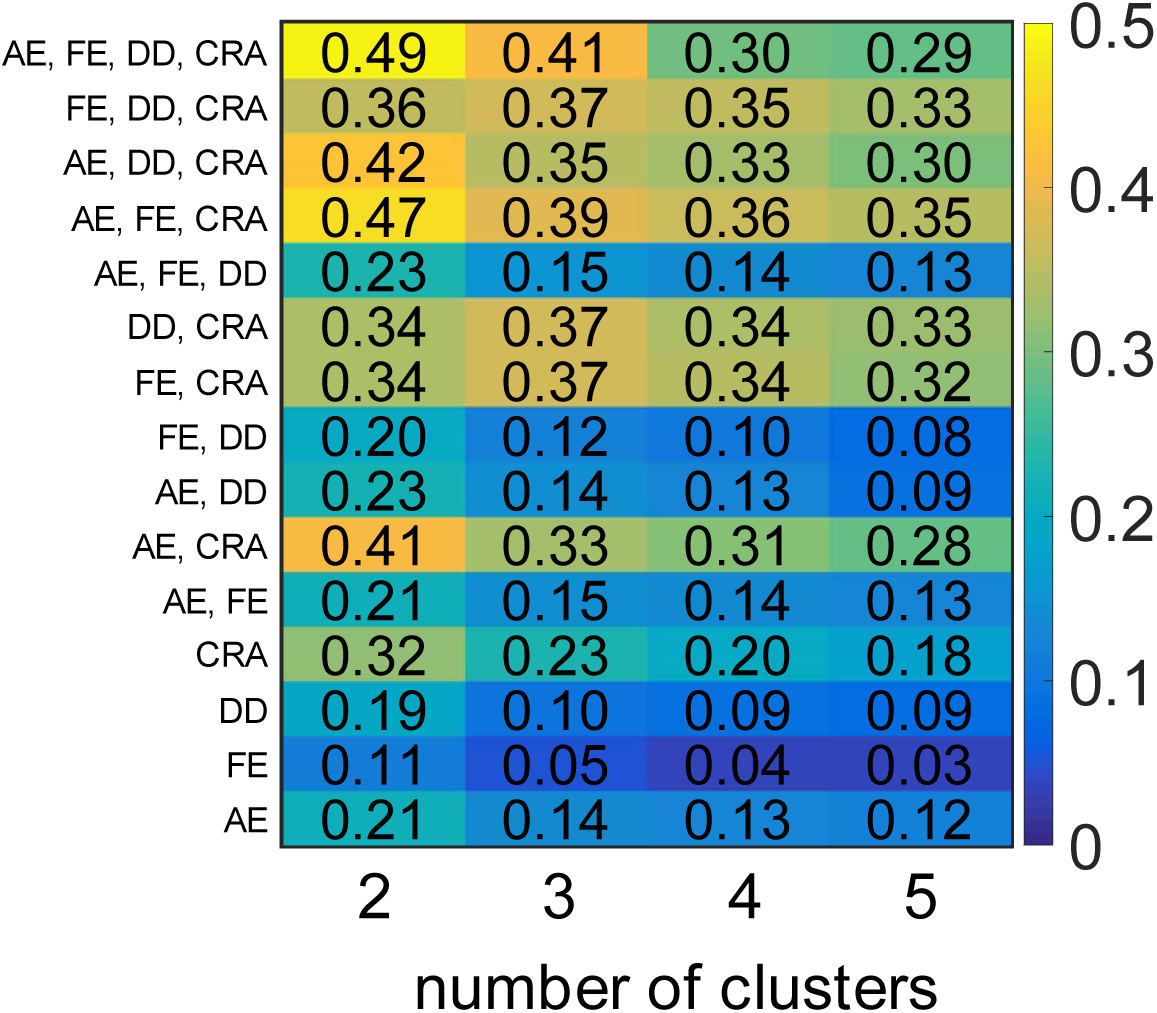
Quality of clustering based on various combinations of decoy-effect indices (noted on the y-axis) and number of clusters (x-axis). Reported are silhouette values for a given number of clusters and combination of decoy-effect indices (AE: attraction effect; FE: frequency effect; DD: dominant vs. dominated; CRA: change in risk aversion). The silhouette can take any values between −1 and 1; higher values indicate that each data point (a measure or set of measures) is closely matched to its own cluster and poorly matched to neighboring clusters. Best clustering results are achieved using all decoy-effect indices and two clusters.

**Supplementary Figure 3.**
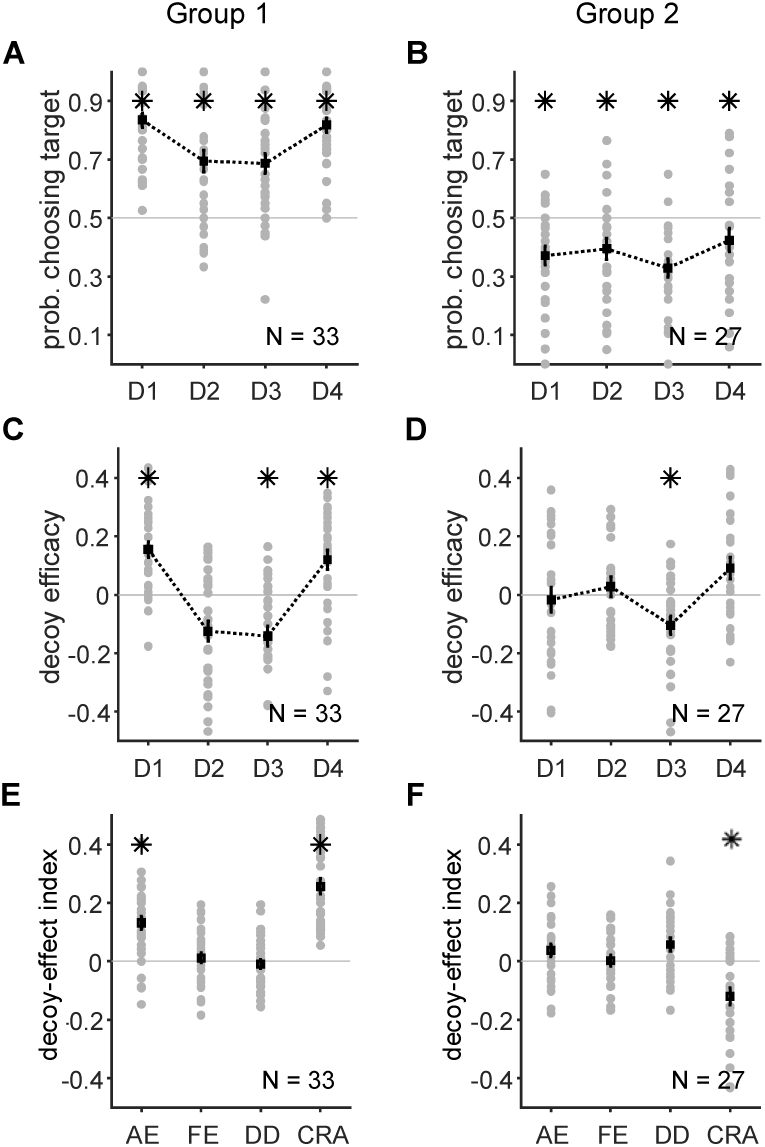
Identified patterns of decoy effects using data from subjects who performed the control condition first and those who performed the control condition only (combined-control data). (**A–B**) Probability of selecting the target for different decoy locations in the two groups of subjects identified by clustering. Each gray circle shows the average probability that an individual subject selected the target for a given decoy location, and black squares indicate the average probability across all subjects. Error bars show the s.e.m., and an asterisk shows that the median of choice probability across subjects for a given decoy location is significantly different from 0.5 (twosided Wilcoxon signed-test, p < 0.05). (**C–D**) Decoy efficacies in the two groups of subjects. Subjects in Group 1 exhibited strong, consistent decoy effects (C), whereas the decoy effects were inconsistent in Group 2 (D). (**E–F**) Plot shows decoy-effect indices for individuals in the two groups of subjects. The first group showed a strong attraction effect and an overall increase in risk aversion. The second group showed a significant decrease in risk aversion.

**Supplementary Figure 4.**
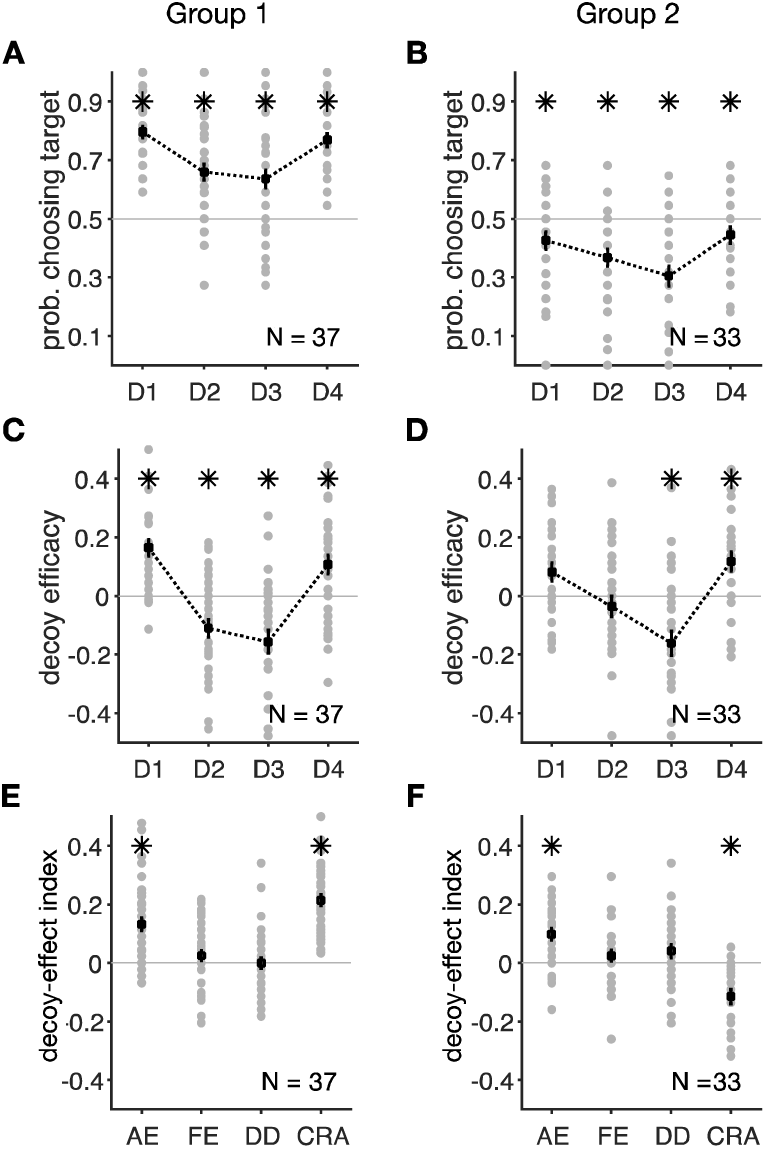
Identified patterns of decoy effects using data from subjects who performed the range-normalization condition first and those who performed the range-normalization condition only (combined-RN data). (**A–B**) Probability of selecting the target for different decoy locations in the two groups of subjects identified by clustering. Each gray circle shows the average probability that an individual subject selected the target for a given decoy location, and black squares indicate the average probability across all subjects. Error bars show the s.e.m., and an asterisk shows that the median of choice probability across subjects for a given decoy location is significantly different from 0.5 (two-sided Wilcoxon signed-test, p < 0.05). (**C–D**) Decoy efficacies in the two groups of subjects. Subjects in Group 1 exhibited strong, consistent decoy effects (C), whereas the decoy effects were inconsistent and limited to decoys next to the less risky gamble (target) in Group 2 (D). (**E–F**) Plot shows decoy-effect indices for individuals in the two groups of subjects. The first group showed a strong attraction effect and an overall increase in risk aversion. The second group showed a significant attraction effect and a decrease in risk aversion.

**Supplementary Figure 5.**
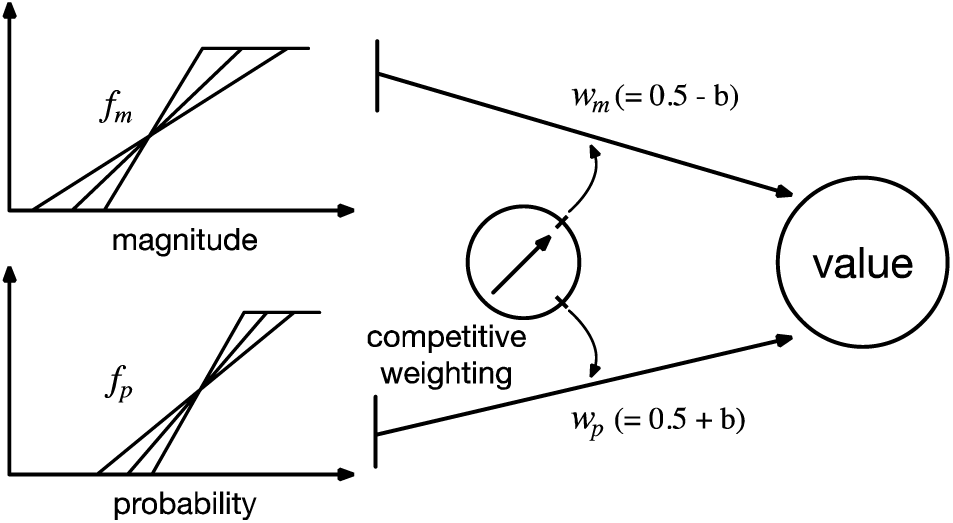
Schematic of the extended model with both low-level and high-level mechanisms. Values of gambles in a given attribute (reward magnitude and probability) are encoded by the corresponding attribute-encoding population of neurons. We assumed that the response of attribute-encoding neurons (response curve) to be a linear function of the attribute value. Representation factor *f*_*i*_ (*i* = {*m, p*}) determines dynamic range (threshold and saturation points) of neural response as well as the slope of the response curve for each attribute. We assumed that an additional competitive mechanism could modulate the weight of each attribute (*w*_*m*_ and *w*_*p*_) on the overall value of each gamble.

**Supplementary Figure 6.**
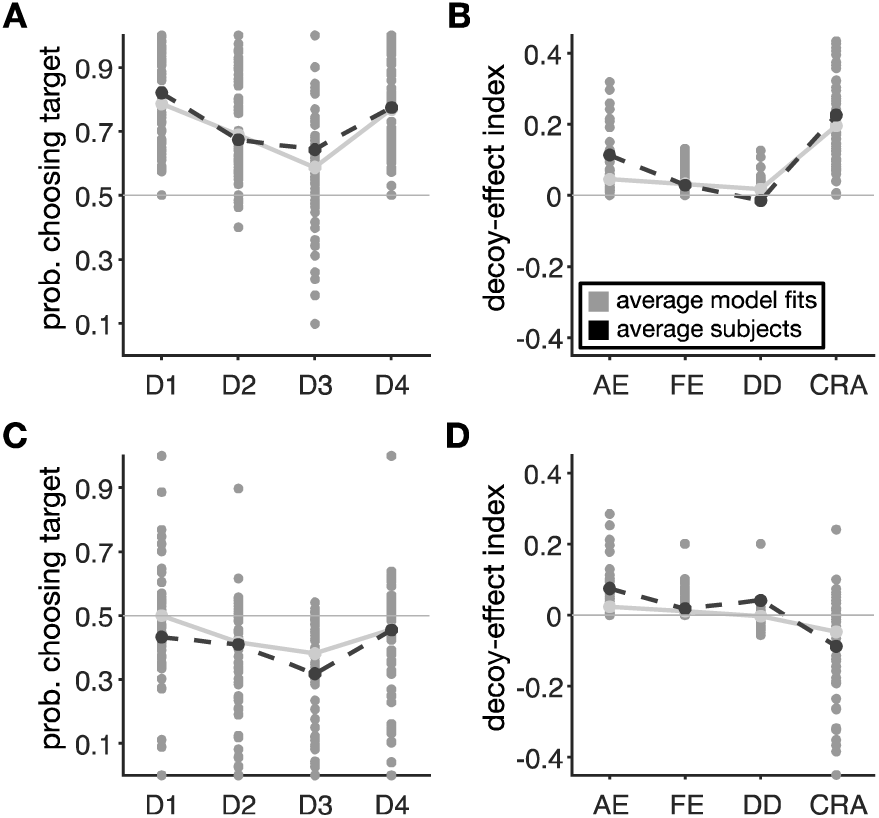
A model with only adjustments of value representations cannot capture all aspects of the experimental data. Plotted are fit of individual subjects based on a model that has adjustments to decoy presentation similar to the original model by Soltani et al. (2012). (**A**, **C**) Each gray circle shows the probability of selecting the target for different decoy locations based on the model fit for individual subjects in the first (A) and second (C) groups of subjects. For comparison, the average value across subjects (black dashed lines) and their fits (gray solid lines) are plotted as well. (**B**, **D**) Plots show four measures for quantifying different effects of decoys on preference based on the cited model fit for individual subjects in the first (B) and second (D) groups of subjects (AE: attraction effect; FE: frequency effect; DD: dominant vs. dominated; and CRA: change in risk aversion).

**Supplementary Figure 7.**
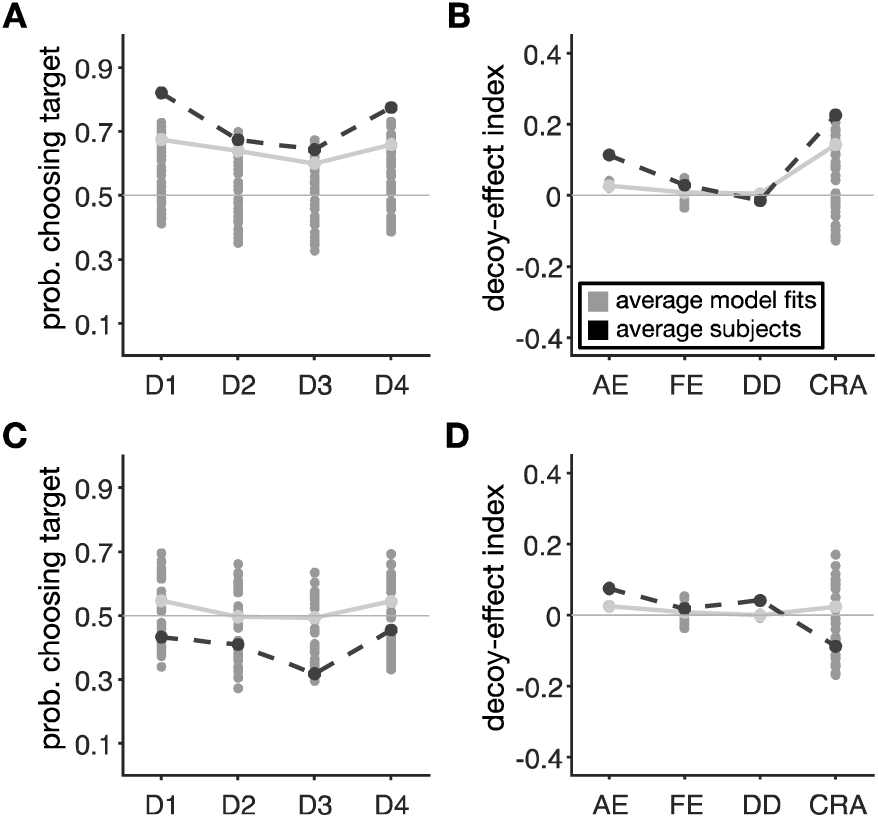
A model with only competitive weighting of reward attributes cannot capture all aspects of the experimental data. Plotted are fit of individual subjects based on a model that only assumes a competitive mechanism to modulate the weight of each attribute (*w*_*m*_ and *w*_*p*_) on the overall value of each gamble. (**A**, **C**) Each gray circle shows the probability of selecting the target for different decoy locations based on the model fit for individual subjects in the first (A) and second groups of subjects. For comparison, the average value across subjects (black dashed line) and their fits (gray solid lines) are plotted as well. (**B**, **D**) Plots show four measures for quantifying different effects of decoys on preference based on the cited model fit for individual subjects in the first (B) and second (D) groups of subjects (AE: attraction effect; FE: frequency effect; DD: dominant vs. dominated; and CRA: change in risk aversion).

**Supplementary Figure 8.**
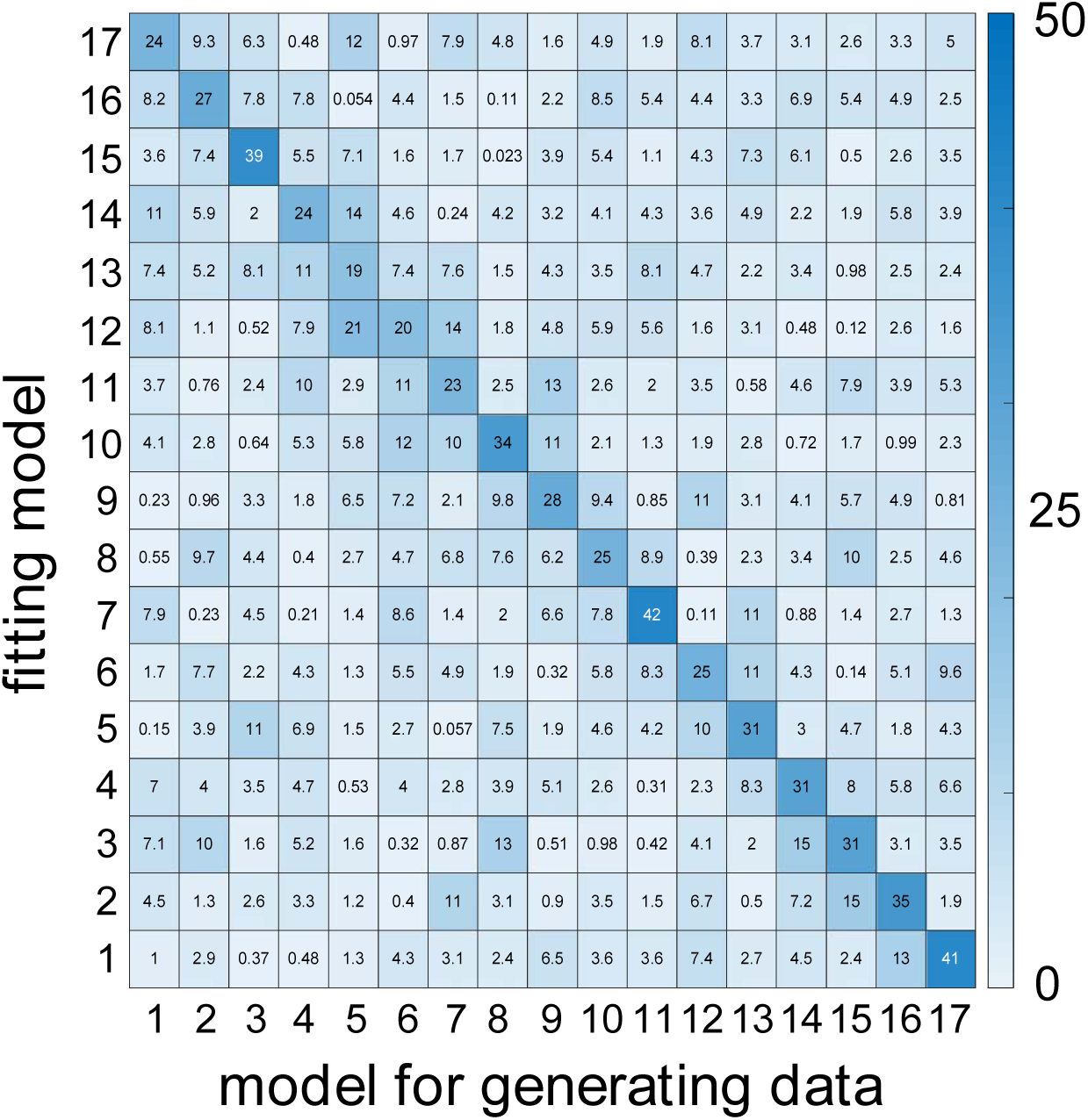
Model recovery. Our fitting method was able to identify the correct model. The value of each cell reports the percentage of instances that a model used to generate the data (shown on the x-axis) was best fit by the same or other models (fitting model, shown on the y-axis. The model corresponding to each number is provided in **Table 1**. The model with the minimum AIC was assigned as the best model. The probability for assigning the best model by chance is ∼6% and thus, values above 19% on the diagonal indicate that in most cases the correct model was identified. For these simulations, we generated 200 sets of data based on a given model and using parameters from the fit of individual subjects’ data, and then fit those data with all the models in order to calculate AIC and determine the best one.

**Supplementary Figure 9.**
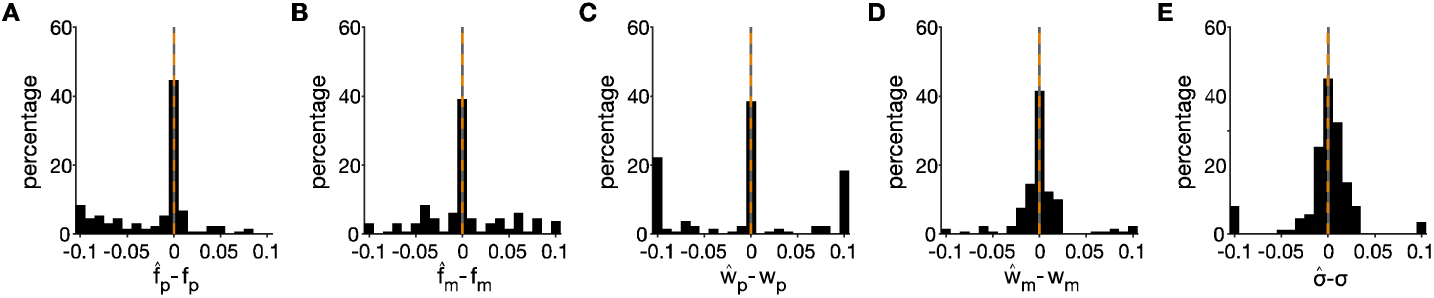
Parameter recovery for the best model. Our method was able to provide unbiased estimates of 5 model parameters. Plots show the difference between estimated and real values for *f*_*p*_ (**A**), *f*_*m*_ (**B**), *w*_*p*_ (**C**), *w*_*m*_ (**D**), and σ (**E**). Orange dashed and solid gray lines show zero and median, respectively. None of the differences are significantly different from zero. Estimation errors are calculated based on 500 sets of randomly generated model parameters.

**Supplementary Figure 10.**
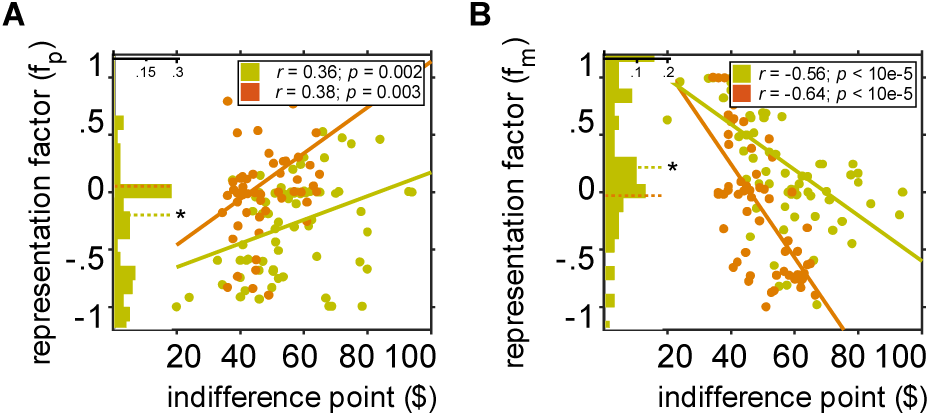
Plots show the estimated representation factor for probability (A) and magnitude (B) as a function of the indifference points within each individual. The green and orange inset histograms plot the fractions of subjects (green: Group 1; orange: Group 2) with certain values of representation factor. Neural representations to decoy presentation of reward attributes were both strongly correlated with the original degree of risk aversion in both groups, reflecting competitive adjustments in how the two reward attributes were processed in the decoy task.

